# MINI-AC: Inference of plant gene regulatory networks using bulk or single-cell accessible chromatin profiles

**DOI:** 10.1101/2023.05.26.542269

**Authors:** Nicolás Manosalva Pérez, Camilla Ferrari, Julia Engelhorn, Thomas Depuydt, Hilde Nelissen, Thomas Hartwig, Klaas Vandepoele

## Abstract

Gene regulatory networks (GRNs) represent the interactions between transcription factors (TF) and their target genes. GRNs control transcriptional programs involved in growth, development and stress responses, ultimately affecting diverse agricultural traits. While recent developments in accessible chromatin (AC) profiling technologies make it possible to identify context-specific regulatory DNA, learning the underlying GRNs remains a major challenge. We developed MINI-AC (Motif-Informed Network Inference based on Accessible Chromatin), a method that combines AC data from bulk or single-cell experiments with TF binding site information to learn GRNs in plants. We benchmarked MINI-AC using bulk AC datasets from different *Arabidopsis thaliana* tissues and showed that it outperforms other methods to identify correct TFs binding sites. In maize, a crop with a complex genome and abundant distal AC regions, MINI-AC successfully inferred leaf GRNs with experimentally confirmed, both proximal and distal, TF-target gene interactions. Furthermore, we showed that both AC regions and footprints are valid alternatives to infer AC-based GRNs with MINI-AC. Finally, we combined MINI-AC predictions from bulk and single-cell AC datasets to identify general and cell-type specific maize leaf regulators. Focusing on C4 metabolism, we identified diverse regulatory interactions in specialized cell types for this photosynthetic pathway. MINI-AC represents a powerful tool for inferring accurate AC-derived GRNs in plants and identifying known and novel candidate regulators, improving our understanding of gene regulation in plants.

## Introduction

Plant development is controlled by a wide variety of endogenous and environmental stimuli that need to be processed correctly to guarantee proper growth along with adequate molecular and physiological responses. One of the main determinants involved in coordinating these different types of signals is the spatiotemporal control of gene expression. Transcriptional gene activity is controlled by transcription factors (TFs), proteins that bind to short and specific DNA sequences, called TF binding sites (TFBS) or motifs, to regulate gene expression (Kulkarni and Vandepoele 2020; Schmitz, Grotewold, and Stam 2022). TFBS are located within cis-regulatory elements (CREs), which are non-coding DNA regions involved in regulating the transcriptional activity of neighboring genes. Although many plant CREs are found within hundreds of base pairs upstream or downstream of the transcription start/end site, they can also be found within introns, untranslated regions (UTRs) and in distal locations from genes (Heyndrickx et al. 2014; Schmitz, Grotewold, and Stam 2022). Distal CREs regulate the expression of target genes (TGs) thousands of base pairs away through chromatin loops that bring them in spatial proximity (Lu et al. 2019). Gene regulatory networks (GRNs) represent a set of regulatory interactions between TFs and their TGs (Mejia-Guerra et al. 2012), and they have been crucial to identify key molecular players and signaling cascades involved in growth, development, and stress responses (Chen et al. 2018; Gaudinier et al. 2018; Reynoso et al. 2019; Jones and Vandepoele 2020; Vercruysse et al. 2021; De Clercq et al. 2021).

Several high- and low-throughput methods exist for the experimental delineation of GRNs (Gaudinier and Brady 2016), starting with techniques that determine the genomic location of TFBS to link them to their putative TGs (TF-based methods). Chromatin immunoprecipitation sequencing (ChIP-seq) profiles the genome-wide TFBS for a TF of interest *in vivo* (Johnson et al. 2007). DNA affinity purification sequencing (DAP-seq) is an *in vitro* alternative to ChIP-seq that enables profiling many TFs in parallel, but that does not consider the epigenetic context of the cells (O’Malley et al. 2016). Protein binding microarrays (PBMs) test the *in vitro* binding of TFs to thousands of short DNA sequences (Franco-Zorrilla et al. 2014). The comparison of genes expressed in wild-type plants with genes expressed in plants overexpressing or defective of a TF makes it possible to identify sets of genes directly or indirectly controlled by the perturbed TF (Krouk et al. 2013). Other approaches are gene-based, such as yeast-one hybrid (Y1H), that test the *in vitro* binding of a specific TF to the promoter of a gene of interest (Meng, Brodsky, and Wolfe 2005). Additionally, there are methods that characterize accessible chromatin (AC) in genomic regions that are depleted of nucleosomes and therefore available for TF binding, potentially controlling transcriptional gene regulation. Prominent AC profiling techniques in plants are Transposase-Accessible Chromatin sequencing (ATAC-seq) (Buenrostro et al. 2015), DNase I hypersensitive sites sequencing (DNase-seq) (Boyle et al. 2008) and micrococcal nuclease sequencing (MNase-seq) (Yuan et al. 2005). MNase-defined cistrome-Occupancy Analysis (MOA-seq) is a modification of the MNAse-seq protocol that increases the resolution of AC regions (ACRs) in the genome by identifying TF footprints, which are small regions (< 30 base-pairs or bps) within AC occluded from DNA cleavage due to TF binding (Savadel et al. 2021).

Most of the previously mentioned experimental assays require laborious and expensive protocols, each one coming with its own limitations, which motivated the advances in computational methods for GRN inference to overcome them (Kulkarni and Vandepoele 2020). The majority of the existing GRN inference methods are expression-based, meaning they merely use expression data to find matching expression profiles of TFs and TGs to predict regulatory relationships (Fu and Medico 2007; Langfelder and Horvath 2008; Huynh-Thu et al. 2010; Roy et al. 2013; Saelens, Cannoodt, and Saeys 2018). While these methods have been successfully used to elucidate regulatory mechanisms (Banf and Rhee 2017; Huang et al. 2018; Haque et al. 2019; Zhou et al. 2020), they suffer from many false positive predictions as no evidence of physical interaction between TF and TG’s regulatory DNA is considered (Gardner and Faith 2005; Marbach, Costello, et al. 2012; Banf and Rhee 2017). The integration of TFBS information can further improve GRN predictions (Marbach, Roy, et al. 2012; Aibar et al. 2017; Ferrari, Manosalva Pérez, and Vandepoele 2022; McCalla et al. 2023), but simply mapping TFBS to the non-coding sequences flanking genes comes with a high rate of false positives. TF motifs are short and degenerate, resulting in low specificity to identify functional TF binding events. To overcome this shortcoming, integrating chromatin accessibility information with TFBS information offers an attractive opportunity to identify more accurate GRNs that reflect the chromatin accessibility landscape of a specific biological sample or condition (Kulkarni, Marc Jones, and Vandepoele 2019; P. Ding et al. 2021). Several methods that integrate different regulatory data types have been developed in recent years. Transcription factor Occupancy prediction By Investigation of ATAC-seq Signal (TOBIAS) is a comprehensive framework that performs TF footprinting on ATAC-seq data and scans the footprints for known motifs (Bentsen et al. 2020). Motif Enrichment in Differential Elements of Accessibility (MEDEA) is a computational tool that identifies cell-type specific accessible regions and performs motif enrichment (Mariani et al. 2020). CellOracle is a machine learning-based tool that integrates single-cell transcriptome and epigenome profiles to infer GRNs (Kamimoto et al. 2023). Integrated regulatory network analysis (IReNA) infers GRN by performing network modularization, transcription factor enrichment, and construction of simplified intermodular regulatory networks (Jiang et al. 2022). The Inferelator 3.0 performs GRN inference by using prior accessibility-based networks that are then improved based on expression data (Gibbs et al. 2021).

Although there are multiple tools available that combine TF motifs with accessibility information, many of these do not perform motif enrichment combined with GRN inference. TOBIAS, for example, identifies TFBS within footprints but does not perform an enrichment analysis nor GRN inference. While MEDEA performs motif enrichment on differential ACRs, it does not infer GRNs and lacks support for plant species. CellOracle, IReNA, and the Inferelator 3.0, integrate TFBS with accessibility information to infer accessibility-based GRNs that are later refined with scRNA-seq data. These accessibility-based GRNs, however, are built by simply scanning motifs in ACRs, resulting in a high number of potential false positives. Furthermore, none of these methods have been evaluated in plants. To fill this gap, we developed MINI-AC (Motif-Informed Network Inference based on Accessible Chromatin), a computational method that integrates TF motif information with bulk or single-cell derived chromatin accessibility data to perform motif enrichment analysis and GRN inference. MINI-AC can be used in two alternative modes - genome-wide and locus-based - to select different non-coding genomic spaces for motif and GRN analysis. We benchmarked MINI-AC in *Arabidopsis thaliana* (herein Arabidopsis) and showed that it outperforms related tools for predicting the correct TFBS for a specific set of ACRs. Next, we implemented MINI-AC for maize (*Zea mays*) and showed that it is able to capture experimentally confirmed proximal and distal regulatory interactions. Finally, we used maize single-cell ATAC-seq data to infer cell-type specific GRNs that helped to identify general and cell-type specific leaf regulators, focusing on C4 metabolism and photosynthesis.

## Results

### Inference of chromatin accessibility-based gene regulatory networks using MINI-AC

MINI-AC is a computational method that integrates TF DNA-binding specificity information with chromatin accessibility data to infer GRNs. MINI-AC’s methodology finds regions in the genome representing potential cis-regulatory sequences known to be bound by specific TFs. Through an efficient TF motif enrichment procedure, MINI-AC predicts regulatory TF-target gene (TG) interactions to generate an accessibility-based GRN (Material and Methods). Each regulon in the GRN, representing a TF and its TGs, is subsequently functionally annotated using Gene Ontology (GO) enrichment analysis, offering insights about TFs controlling specific biological processes (Figure 1A).

**Figure 1.**
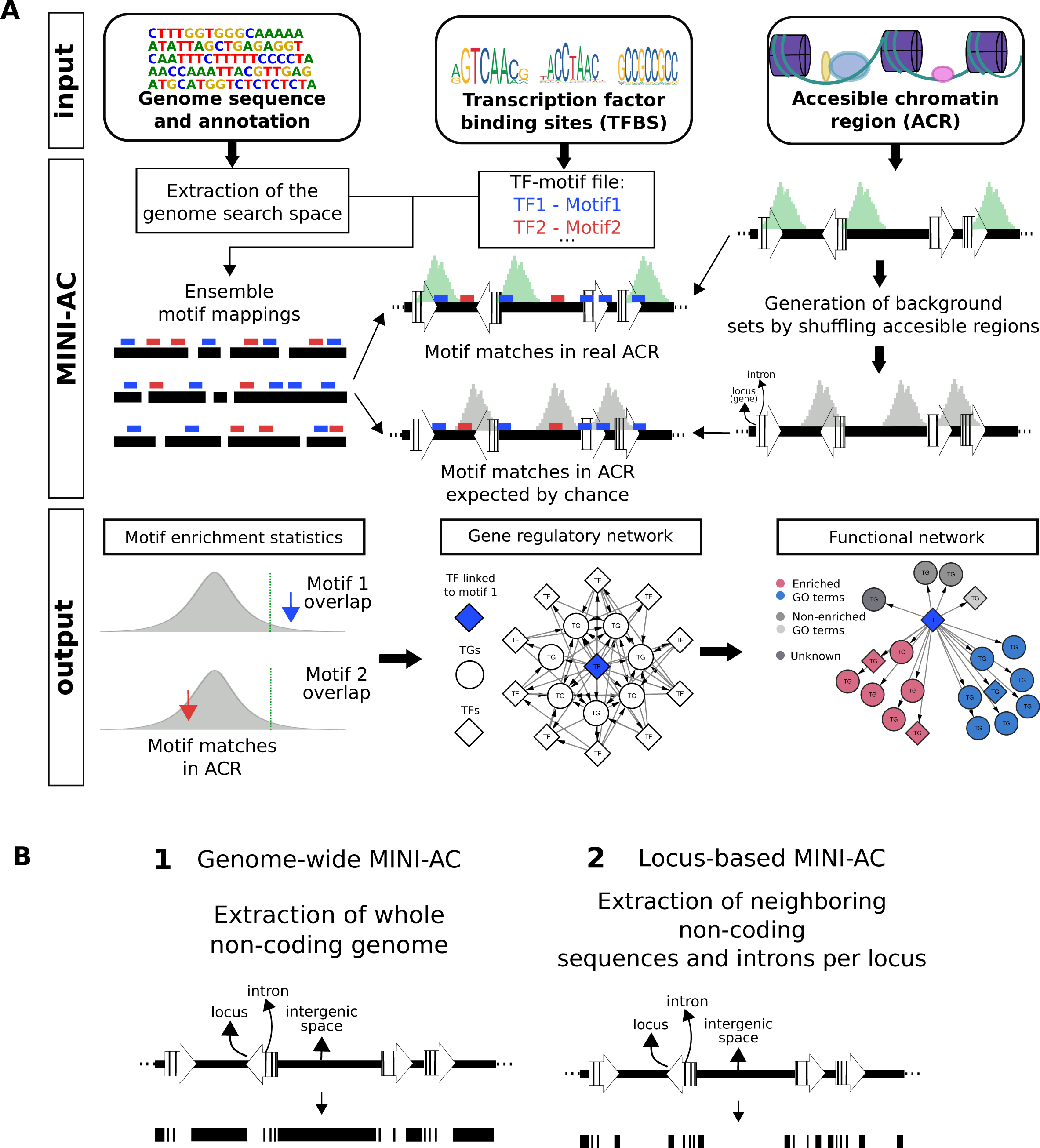
Overview of MINI-AC and its modes. (A) Overview of MINI-AC methodology, inputs and outputs. MINI-AC receives as input a genome sequence and annotation, TFBS information, and a user-provided ACR file. As output, it generates information about motifs showing enrichment on the ACRs, a network that is context-specific for the provided ACRs and a functional enrichment analysis of the regulons in the GRN. (B) Overview of the two modes implemented in MINI-AC, depending on the non-coding genomic search space used for motif mapping: genome-wide and locus-based.

MINI-AC has three main inputs: a sequenced and annotated genome, motifs representing known TFBS and chromatin accessibility data represented as a set of ACRs or footprints. The first step of MINI-AC is the extraction of the non-coding genomic search space and the mapping of known TF motifs to obtain motif mappings, which are a set of genomic locations with putative TFBS, referred to as motif matches. This step of the pipeline is pre-computed per genome. In a second step, the motif mappings are used to identify enriched motifs in a set of ACRs. The enriched motifs are identified by generating a background model of the input ACRs to estimate the number of motif matches in ACRs expected by chance, and comparing them with the real number of motif matches in ACRs. The TFs associated with motifs that show a significant enrichment in the ACRs are linked to TGs, based on proximity, to define regulons (Materials and Methods and https://github.com/VIB-PSB/MINI-AC). The motif enrichment statistics (π-value, a metric that combines q-value and enrichment fold; Materials and Methods) are used to rank motifs and the associated TFs.

MINI-AC is available as a Nextflow (Di Tommaso et al. 2017) pipeline and contains pre-computed motif mappings for Arabidopsis (Araport11) and maize (AGPv4). MINI-AC can be run in two different modes, genome-wide and locus-based, depending on the non-coding genomic space that is used to perform the motif mapping. In the genome-wide mode, the motifs are mapped to the whole non-coding genome, while in the locus-based mode, the motifs are mapped to the flanking (upstream and downstream) non-coding and intron sequences of each locus in the genome (Figure 1B). While the genome-wide mode is expected to capture more ACRs, the locus-based is expected to have a denser motif signal, given that the majority of the TFBSs are less than 2kb from genes (Heyndrickx et al. 2014; Tu et al. 2020), and this influences the final GRN composition. For example, the locus-based mode with 5 kb upstream of the translation start site (TrSS), introns, and 1 kb downstream of the translation end site (TrES) covers 73.5% of the Arabidopsis non-coding genome, but only 9.6% of the maize non-coding genome (Figure S1). This discrepancy is due to the size differences in intergenic space: the median is 1,220 bps for Arabidopsis and 23,714 bps for maize.

The motif mappings are a crucial component of MINI-AC, as their quality determines the completeness and accuracy of the inferred GRNs. Previously, we have shown that the motif mapping tools Find Individual Motif Occurrences (FIMO) and Cluster-Buster (CB) generate complementary motif matches and that combining matches from FIMO with top-scoring matches from CB in an ensemble approach gives better results to recover ChIP-based TF binding events, compared with using a single motif mapping tool (Kulkarni, Marc Jones, and Vandepoele 2019). However, using all motif matches identified by FIMO or CB gives an excessive number of candidate target genes for some motifs and interferes with successfully applying motif enrichment. Therefore, selecting a number of top-scoring motif matches used for GRN inference is an important factor when running MINI-AC. We calibrated an optimal number of top-scoring motif matches for the two motif mapping tools using ChIP-based gold standard sets of TFBS (Methods S1). For each motif mapping tool and each species, we evaluated the number of top-scoring motif matches found within the ChIP-seq peaks of the corresponding TF to compute precision, recall, and F1 (Figure S2). In Arabidopsis, for FIMO and CB the optimal top scoring matches (highest F1 value) were found at top 7,000 and top 4,000, respectively. For maize, the FIMO optimal threshold was found at 16,000 and the CB optimal top at 8,000. These top-scoring motif matches were combined into ensemble motif mappings for each species, containing motifs for 1117 and 1234 TFs for Arabidopsis and maize, respectively.

Next, we assessed whether the ensemble motif mappings for maize yielded increased performance to detect functional TFBS, as previously shown in Arabidopsis (Kulkarni, Marc Jones, and Vandepoele 2019), by evaluating motif mapping results of FIMO and CB using ChIP-seq peaks for 68 TFs (Methods S1). Scoring the unique and shared motif matches of each motif mapping tool on these ChIP-seq peaks (i.e. peaks with a motif match of the profiled TF) revealed that combining the 16,000 top-scoring FIMO motif matches with the 8,000 top-scoring CB motif matches in maize resulted in an 15% increase (from 63% using only FIMO to 78% using both) of the number of ChIP-seq peaks that contained a correct TF motif match, compared with using only the top 16,000 FIMO motif matches. Therefore, these results indicate that the FIMO and CB complementarity also applies for maize.

To generate GRNs using the genome-wide mode, the motif matches within ACRs are annotated to the closest gene, but in cases where the distance to the flanking closest genes is similar, it is uncertain that the closest gene is the correct TG. We evaluated, for cell-type specific ACRs, in how many cases the second-closest gene was differentially expressed (DE) in that specific cell-type compared to the closest gene (Materials and Methods and Methods S2). We used cell-type specific ACRs from maize mesophyll and bundle sheath (Materials and Methods), annotated them to the two closest genes, and then measured the proportion of closest and second-closest genes that were DE in each cell type. This analysis revealed that, although annotating the two closest genes increased the total number of DE genes compared to annotating only the closest gene (Figure S3), this also resulted in a decrease in the ratio of DE genes vs. non-DE genes (from 0.074 up to 0.052 in mesophyll and from 0.05 up to 0.038 in bundle sheath). This resulted in a decrease in precision, and a F1 decrease of up to 0.026 in mesophyll and 0.018 in bundle sheath. Therefore, by default genome-wide MINI-AC annotates only the closest TG that can be located far up- or downstream of the ACR, but this can be changed in the MINI-AC parameter settings. In conclusion, we present MINI-AC as a motif enrichment and GRN inference method implemented and optimized for Arabidopsis and maize.

### MINI-AC outperforms alternative motif enrichment tools in predicting cell-type specific regulators in Arabidopsis

To evaluate the ability of MINI-AC to identify functional TFBS and learn accurate GRNs, we benchmarked the motif enrichment method implemented in MINI-AC against two other tools that also test genomic interval’s enrichment: Giggle (Layer et al. 2018) and Bedtools Fisher (Quinlan and Hall 2010). We selected them because they can be run for any species, and both tools use a different enrichment approach compared to MINI-AC. They estimate the significance of overlap and enrichment between two sets of genomic intervals by computing the overlapping and unique regions in each set and applying a Fisher’s Exact two-tailed test. In this study, the two sets of genomic intervals are ACRs and motif mappings. While Giggle speeds up the computation using a web search engine algorithm for many files at once, Bedtools Fisher performs enrichment for one pair of files at a time. Both tools, and MINI-AC, generate motif enrichment statistics, which include a q-value (p-value corrected for multiple hypothesis testing) that reflects the statistical significance of the overrepresentation of a motif within the ACRs, and a fold enrichment, reporting how much bigger this overrepresentation is than expected by chance.

To compare these tools with MINI-AC, we evaluated how many cell-type specific DE TFs each tool could predict given a set of cell-type specific ACRs. Thus, we selected three different datasets containing bulk cell-type specific ACRs from Arabidopsis stem cells, phloem cells, and epidermis cells (Materials and Methods). These datasets were selected because they report cell-type specific peaks (RB and S 2011) and they had a paired RNA-seq dataset, or there was expression information available for those cell types, that can be used to determine if the TF associated with the enriched motifs is DE in the respective cell type (Methods S2). We also included a fourth dataset comprising “synthetic” ACRs made by combining peaks of 19 ChIP-seq-profiled TFs overlapped with ACRs from Arabidopsis seedlings. The goal of this synthetic dataset is to have a set of ACRs filtered for specific TFBS expected to, ideally, be enriched and recovered by the different enrichment tools. This is in contrast to the three other datasets, where the expected number of binding TFs showing motif enrichment is unknown and estimated using the paired gene expression information.

To perform a fair comparison, we first determined for each tool the q-value cutoff that controls false positives in an equally stringent manner (Materials and Methods). Next, we processed the four datasets with each tool, applying the q-value cut-offs at false discovery rate (FDR) 0%, and compared the results. We found that MINI-AC (either locus-based or genome-wide) had the highest recall, which is the proportion of the correct motifs each method predicted as enriched, and highest precision (except for stem), which is the proportion of predicted enriched motifs that are correct. This resulted in MINI-AC having the best overall performance (highest F1 value; Figure 2A), except for the “synthetic” ACR dataset, for which Giggle yielded the highest F1 by a small margin (0.088 for Giggle, 0.066 for Bedtools, 0.067 for genome-wide MINI-AC, and 0.068 for locus-based MINI-AC). For the rest, MINI-AC’s performance was better, especially for the ACR datasets of epidermis, phloem, and stem, where Giggle and Bedtools showed a very low recall. For the “synthetic” ACR dataset, the recall of Giggle and Bedtools showed a high improvement (0.855 and 0.600 respectively), yet unable to match MINI-AC’s (0.973 on average). This improved recall in the “synthetic” dataset (only 19 TFs) could be due to a stronger and less complex motif signal compared to bulk ACR datasets, as in the latter probably hundreds of different TFs are active. For the epidermis ACR dataset, Giggle and Bedtools failed to retrieve any of the motifs associated to DE TFs in that cell type. This result could be due to the low number of motifs associated with DE TFs in stem (only 29, while other cell types have between 150 and 182), as well as the low number of ACRs (Table S1). When comparing the two different MINI-AC modes, their performance in precision and recall was very similar, probably because the non-coding genomic space defined for the locus-based MINI-AC mode (5 kb upstream of the TrSS, 1 kb downstream of the TrES, and introns) covers most of the non-coding genome of Arabidopsis that is used in the genome-wide mode (73.5%).

**Figure 2.**
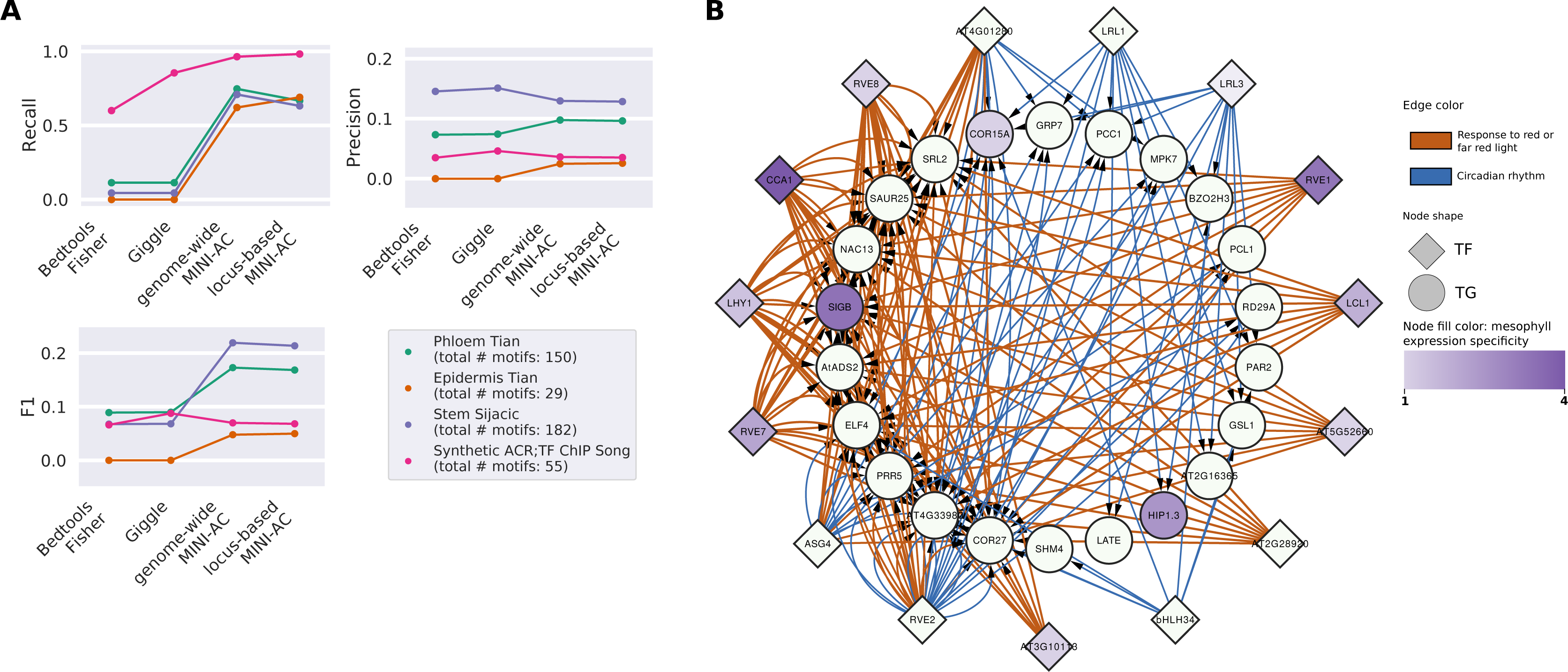
Benchmark of MINI-AC in Arabidopsis. (A) Comparison of performance statistics (recall, precision, and F1) for Bedtools Fisher, Giggle, and genome-wide and locus-based MINI-AC modes. For each dataset used in the comparison, the legend indicates the number of motifs expected to be retrieved as enriched by the different tools (true positive motifs). (B) Network showing the regulatory interactions predicted by MINI-AC for mesophyll that show an enrichment for GO annotations (q-value < 0.01) of “response to red or far red light” (in red) and “circadian rhythm” (in blue). The TFs shown are regulators predicted to control the expression of TGs enriched for the mentioned GO terms. The edge width represents the -log10(q-value) of GO enrichment, while the node color represents the mesophyll expression specificity, measured as -log2(mesophyll expression rank / stem expression rank).

Besides assessing MINI-AC’s motif enrichment performance, we also evaluated the quality of the GRNs inferred based on the motif enrichment results. We used a mesophyll cell-type specific ACR dataset from (Sijacic et al. 2018), which was not used in the benchmark, to infer GRNs with MINI-AC (Giggle and Bedtools were excluded from this analysis because they are not GRN inference methods). This dataset contains differential ACRs of mesophyll cells compared to stem cells. Therefore, to evaluate the predicted regulators and TGs, we used genes that were strongly expressed in mesophyll compared with stem cells (Methods S2). MINI-AC predicted 632 enriched motifs that are associated with 552 TFs (Data S1). The inferred GRN reports 83,419 interactions and 1533 TGs. Out of the 158 TFs showing mesophyll-specific expression and expected to be associated with enriched motifs, 68% (108) were present in the MINI-AC network. Functional analysis of the regulons in the MINI-AC mesophyll GRN revealed enrichment for processes relevant for leaf and mesophyll function, such as “circadian rhythm” (57 TFs) and other light-related processes, mainly “response to red or far red light”. We filtered the mesophyll network for regulons showing enrichment for these two GO terms and verified if known regulators of circadian rhythm and light perception in leaf were found (Figure 2B). Among the TFs predicted by MINI-AC to regulate response to red light and circadian rhythm, CCA1, LHY1, RVE1/2/6/7/8, and LHY/CCA1-like 1 were found, which have been previously reported as TFs controlling these processes (Creux and Harmer 2019). Additionally, some of these TFs, such as CCA1, or RVE1 show a strong mesophyll expression specificity (large difference in expression ranking compared to stem cells; Methods S2). So MINI-AC not only predicts relevant TFs known for mesophyll, but also links them to TGs involved in the biological processes controlled by these TFs. On the other hand, MINI-AC also reports functional enrichment for regulons controlled by unknown TFs, such as “response to red or far red light” for AT3G10113, indicating that MINI-AC can be used to generate new testable hypotheses for specific regulators.

Overall, these results indicate that, compared with other tools that perform enrichment between two sets of genomic intervals, MINI-AC shows a better performance in predicting motif enrichment of DE TFs in cell-type specific ACRs. Next, our results suggest that MINI-AC performs well on ACR datasets with high complexity (potentially containing hundreds of TFBS). Finally, MINI-AC predicted the correct functional enrichment for the TGs of known regulators involved in light perception and circadian rhythm in an Arabidopsis mesophyll GRN, highlighting its potential to identify novel regulators.

### Prediction of proximal and distal regulatory interactions from bulk ACRs in maize

Maize is a crop and model species of high economical relevance, yet the inference of GRNs remains challenging for this species due to the large intergenic regions and the presence of distal regulatory elements (Lu et al. 2019). We evaluated if MINI-AC can infer accurate GRNs in maize that capture proximal and distal regulatory interactions. Different MINI-AC versions for maize were implemented, either using the genome-wide mode or three non-coding genomic spaces of the locus-based mode, small (1kb upstream of the TrSS, introns, and 1 kb downstream of the TrES), medium (5 kb upstream of the TrSS, introns, and 1 kb downstream of the TrES), and large (15 kb upstream of the TrSS, introns, and 2.5 kb downstream of the TrES). For each version, we evaluated how well it could predict (1) enrichment of motif/TFs and (2) relevant TF-TG interactions. A maize leaf bulk ATAC-seq ACR dataset (32,481 ACRs covering 13 Mbps) (Lu et al. 2019) was used as input, and these results were evaluated using a gold standard of ChIP-seq-derived TF-TG interactions in maize mesophyll for 104 TFs (Tu et al. 2020), of which 62 have one or more associated motifs. To evaluate the MINI-AC networks, the gold standard was adapted to each MINI-AC mode and non-coding genomic definition, so that a theoretical recall of 100% could be obtained. We did this by selecting the TFs with available motif information, annotating only the peaks that contained the motif associated with the incoming TF, and discarding the peaks outside the corresponding non-coding genomic region definition. The sizes of the gold standards adapted to the different non-coding genomic region definitions are summarized in Table S2.

First, we evaluated the performance of MINI-AC in predicting enriched TFs (62) and motifs (489) present in the gold standard (Figure 3A). The small and medium locus-based modes had the highest TF retrieval rate (both with a recall of 0.903 compared with 0.871 for genome-wide). The genome-wide mode had the highest precision (0.054) and F1 value (0.102) compared with the three locus-based modes (F1 value ranging from 0.096 to 0.100). Next, we computed the precision-recall curve and the area under the precision-recall curve (AUPRC) to measure the ability to predict correct motifs while minimizing incorrect ones. If the motifs linked with gold standard TFs have low ranks in the MINI-AC enrichment, the AUPRC will be higher with high precision at low ranks. The results showed that the four tested MINI-AC versions had a very similar AUPRC (between 0.425 and 0.449; Figure 3C and Figure S4), with locus-based small showing the highest AUPRC by a small margin compared to locus-based medium (0.449 and 0.446 respectively).

**Figure 3.**
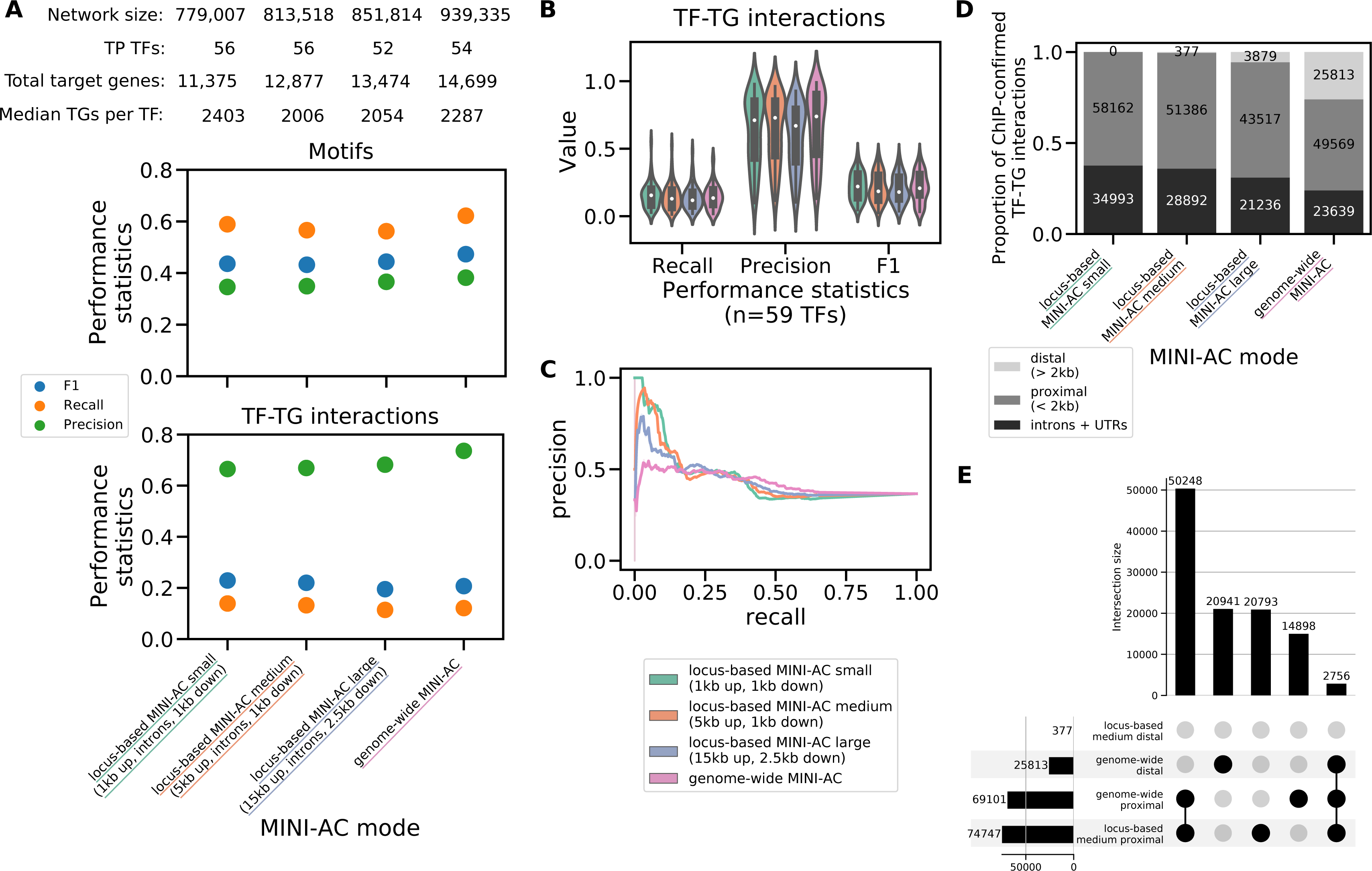
Performance comparison of MINI-AC modes at predicting proximal and distal regulatory interactions using a leaf maize bulk dataset. (A) Comparison of performance statistics in predicting maize leaf GRNs for the three non-coding genomic spaces in locus-based MINI-AC, and genome-wide MINI-AC. The plots show the performance statistics in predicting enrichment for motifs associated with TFs from the gold standard (upper plot) and TF-TG interactions from the gold standard (lower plot). (B) Violin plot showing the distribution of performance statistics for all the TFs in the gold standard individually. (C) Precision-recall curve of retrieval of enriched motifs by MINI-AC that are associated with TFs of the maize leaf gold standard, for the different methods tested (locus-based or genome-wide). (D) Stacked barplot showing the proportion of proximal (< 2kb from the closest gene), genic (introns + UTRs), and distal (> 2kb from the closest gene) ChIP-seq confirmed interactions predicted by each method. (E) UpSet plot showing the number of unique and shared ChIP-confirmed regulatory interactions (proximal and distal) between the genome-wide and locus-based (medium non-coding genomic search space) MINI-AC modes.

Second, to evaluate the MINI-AC GRNs for each mode, we compared the inferred TF-TG interactions with the TF-TG interactions of the corresponding gold standard (Figure 3A and Figure 3B). The genome-wide mode produced the GRN with the highest precision (0.736) of ChIP-confirmed TF-TG interactions. However, the locus-based small mode predicted the highest fraction of gold standard interactions (0.139 recall). The final F1 values for TF-TG interaction predictions, from highest to lowest, were as follows: locus-based small (0.229), locus-based medium (0.220), genome-wide (0.195), and locus-based large (0.195) (Figure 3A).

While these different comparisons revealed no major differences in the global recovery of known regulatory interactions using the AC-derived GRNs, we next assessed if the genome-wide MINI-AC mode is more successful than the locus-based in inferring distal regulatory interactions. For each MINI-AC mode and non-coding genomic region definition, we labeled each correct TF-TG interaction into 3 classes: “introns + UTR”, “proximal” (< 2kb from the closest gene), and “distal” (> 2kb from the closest gene) (Figure 3D). Comparing the different classes revealed that genome-wide MINI-AC predicts 28% of its correct TF-TG interactions as distal, while medium and large locus-based MINI-AC mode predict 0.5% and 6%, respectively. When considering only the correct interactions predicted uniquely by genome-wide MINI-AC, this percentage increases to 50%, meaning that half of the correct interactions retrieved uniquely by genome-wide MINI-AC are distal. Finally, we evaluated if the correct distal and proximal TF-TG interactions found by the genome-wide MINI-AC were unique or shared with the medium locus-based mode (the best performing locus-based version that can predict distal interactions based on F1 score). We observed that approximately 81% (21,000) of the distal TF-TG interactions found by genome-wide MINI-AC mode were unique, and not found by locus-based mode neither proximally nor distally (Figure 3E). On the other hand, the locus-based mode retrieved 29% (20,941) of unique proximal interactions, while the genome-wide mode retrieved 22% (20,793).

While ATAC-seq provides ACRs that span a few hundreds of DNA base pairs, it lacks resolution about the exact location of TFBS. To overcome this, experimental and computational methods have been developed to identify TF footprints. To assess if MINI-AC can infer motif enrichment and GRNs from footprints, in contrast to larger ATAC-seq peak regions, we generated a set of genomic footprints and peak regions applying MOA-seq in maize leaf and used them as input for the genome-wide mode (Materials and Methods). We evaluated the motif enrichment performance by using a combined set of DE genes of different leaf cell-types (Methods S2). Conversely, to evaluate the inferred GRNs, we compared them to the previously mentioned ChIP-seq gold standard. Regarding motif enrichment, the peaks and the footprints datasets yielded 812 and 719 enriched motifs, respectively, of which 617 were shared between the two. The results, summarized in Table S3, show that both data types (peaks and footprints) showed similar performances when retrieving correct motifs as enriched, with the peaks showing better F1 (0.580 for peaks vs. 0.529 for footprints). While the precision-recall curve revealed that MOA-seq footprints are superior to identify correct TFs among the top ranked motifs, the AUPRC showed a better overall performance when using MOA-seq peaks (AUPRC for peaks 0.576 and AUPRC for footprints 0.547; Figure S5A and S5B). For GRN predictions, both dataset types generated networks with similar sizes (4,110,921 interactions using peaks and 3,560,086 interactions using footprints) while the evaluation of ChIP-seq confirmed interactions revealed that peaks yielded a better F1 (0.275) in comparison with the GRN predicted using footprints (F1 of 0.242). A comparison of the correct TF-TG interactions of each network showed that there was a large overlap between the correct edges of the GRNs (Figure S5C-D; Jaccard similarity coefficient of 0.653).

Overall, these results revealed that MINI-AC can be successfully employed to infer GRNs in maize. While the genome-wide mode retrieved more motifs from the gold standard TFs as enriched and predicted the most precise GRNs, the locus-based small mode showed better prioritization of TFs and retrieved the most complete GRNs. Additionally, the GRNs predicted by the genome-wide mode contained four times more ChIP-confirmed distal regulatory interactions than the locus-based mode’s GRNs, showing it is the most appropriate option to use with species like maize with large intergenic spaces and many distal regulatory elements. Finally, we demonstrate that using MOA-seq (peaks and footprints) provides a valid alternative to ATAC-seq as input for MINI-AC.

### Leveraging MINI-AC predictions from bulk and single-cell AC datasets highlights general and specific leaf regulators

The leaf is the main organ where photosynthesis occurs, producing energy and carbohydrates that sustain the plant’s growth and life cycle (Vercruysse et al. 2021). In recent years, different maize leaf bulk datasets have been published (Dong et al. 2017; Lu et al. 2019; Tu et al. 2020), providing ACRs for different leaf cell types combined, which is in contrast with single-cell technologies that profile accessibility landscapes of individual cells and cell types (Angerer et al. 2017; Marand et al. 2021). Marand and co-workers (Marand et al. 2021) profiled six maize organs (among them seedling, that contains leaf cells) using single-cell ATAC-seq (scATAC-seq). Our first goal was to evaluate if MINI-AC can infer GRNs from single-cell-derived data, and next compare these results with MINI-AC predictions using bulk datasets. We performed motif enrichment and GRN inference with MINI-AC using 3 publicly available maize leaf bulk ATAC-seq datasets (Dong et al. 2017; Lu et al. 2019; Tu et al. 2020), one leaf bulk MOA-seq dataset (Materials and Methods), and the single-cell dataset from Marand et al. (Marand et al. 2021) (referred to as “Marand dataset”). The Marand dataset was subsampled for 10 leaf cell types, of which we selected the most specific peaks (using z-score specificity; Materials and Methods), that has on average 39% of distal peaks for all the cell types, in comparison to 33% in the bulk datasets. We selected the genome-wide MINI-AC mode, as these distal peaks would be discarded when using the locus-based mode (Figure S6A-C) and we observed that distal peaks tend to be more cell-type specific (Figure S6D). The bulk- and single-cell-derived ACR sets were used for motif enrichment and GRNs inference, but to focus on the functional regulatory interactions in leaf, the predictions were filtered for leaf-expressed genes (Methods S3).

To test MINI-AC’s ability to use single-cell derived datasets, we evaluated the motif enrichment results of the Marand dataset cell types by combining published datasets of single-cell or single-nuclei-derived RNA-seq DE genes from different leaf cell types: mesophyll, bundle sheath, guard cell, subsidiary, and pavement cell (Methods S2). Additionally, for each of these cell-types, we generated a functional GRN (Data S2). We found that 30%, 29%, and 34% of the motifs predicted as enriched are recognized by DE TFs in mesophyll, subsidiary, and bundle sheath, respectively. These numbers increased to 44%, 35%, and 36% when considering only the top 150 enriched motifs (Table S4), showing that prioritizing motifs by their significance can improve the retrieval of cell-types specific DE TFs. For guard cell, given the low number of DE genes and TFs, the percentage of motifs associated with DE TFs was very low (3%).

Next, we compared the enrichment results using singe-cell-derived and bulk-derived ACRs, and analyzed the total enriched motifs and DE enriched motifs (Figure S7A and S7B). We observed a large agreement between both ACR data types, so we resorted to differences in the motif enrichment ranks to identify TFs showing specificity. We compared the rank of the DE TFs that were within the top 150 ranked motifs in any of the datasets, and used Borda rank (geometric mean of all ranks) to identify TFs with a consistent low rank across all datasets (high confidence regulators) and TFs with low rank in one cell-type compared to other datasets (cell-type specific regulators). Gene identifiers for the different genes reported below can be found in Table S5.

To find putative general leaf regulators, we focused on the top 150 ranked TFs showing differential expression with the lowest Borda ranks (Figure 4 and Table S6). The TF with the lowest Borda rank across all bulk and single-cell datasets was *Zm00001d006034*, an ortholog of Arabidopsis AHL10 (*AT-HOOK MOTIF NUCLEAR LOCALIZED PROTEIN 10*) involved in growth regulation during stress in Arabidopsis (Wong et al. 2019), but which has not been characterized in leaf development. Other TFs showing consistently low ranks were HB78 (up-regulated in subsidiary), HB62 (down-regulated in mesophyll), and HB11, which showed the highest motif enrichment in mesophyll and subsidiary (rank 3 and 4 respectively; same rank because they are associated with the same motif). HB78 is an ortholog of Arabidopsis HAT3, shown to have a critical role in establishing the dorso-ventral axis in cotyledons and developing leaves (Bou-Torrent et al. 2012), while HB62 is an ortholog of Arabidopsis HAT22, which has been reported to function in leaf development and act as a repressor of an osmotic stress related regulator (Preciado, Begcy, and Liu 2022; Seok et al. 2022). The gibberellic acid (GA) mediated signaling pathway was among the enriched functional categories for the TGs of HB62 and AHL10 in mesophyll. GA plays a key role in leaf development and size of the division zone (Strable and Nelissen 2021). Among those TGs there is gibberellin 20-oxidase1 (GA20OX1), which is determinant of the division zone size (Nelissen et al. 2012).

**Figure 4.**
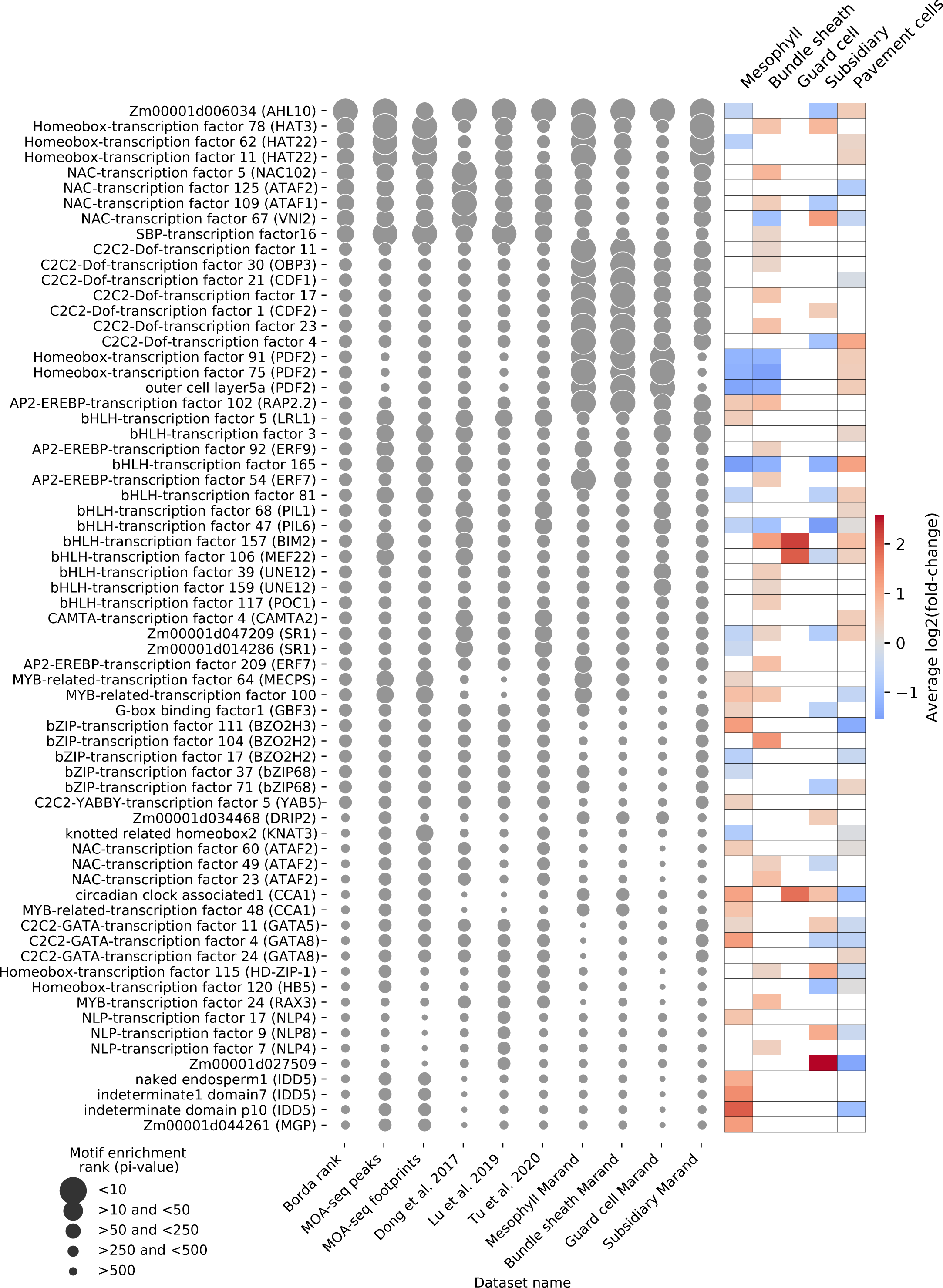
Motif enrichment rank comparison of cell-type specific DE TFs in leaf single-cell and bulk datasets in maize. Bubble plot showing the ranks of all DE TFs associated with motifs within the top 150 enriched motifs in any dataset (bulk or single-cell ACR sets). The rows are annotated based on the up- or down-regulation status of the TF in mesophyll, bundle sheath, guard cell, subsidiary or pavement cell (red for up-regulated genes and blue for down-regulated genes). The blue or red shade is proportional to -log2(fold change), averaged for cell types shared between the single-cell and single-nuclei RNA-seq datasets used. The bubble size represents the motif enrichment rank, with larger bubbles indicating lower ranks.

To find putative cell-type specific regulators, we focused on TFs with low ranks in specific cell-types (Data S3). Based on 79 cell-type specific DE TFs in mesophyll, 54 (68%) were predicted by MINI-AC as enriched in the mesophyll specific ACRs. The DE TFs EREB102 and EREB54 showed high enrichment in mesophyll compared to other cell-types. EREB102 was up-regulated in mesophyll and its regulon showed high enrichment for mesophyll up-regulated genes (17% of TGs), suggesting that EREB102 is an important mesophyll regulator. The regulon predicted by MINI-AC for this TF in mesophyll was functionally enriched for photosynthesis, chloroplast organization and response to light. MYBR100 also showed specificity for mesophyll (Figure 4 and Figure 5) and is an ortholog of Arabidopsis RVE6 (*REVEILLE6*), involved in the regulation of circadian rhythm and mesophyll cell size determination (Gray et al. 2017). The MYBR100 regulon showed functional enrichment for “circadian rhythm” and enrichment for mesophyll up-regulated genes (18% of DE TGs).

**Figure 5.**
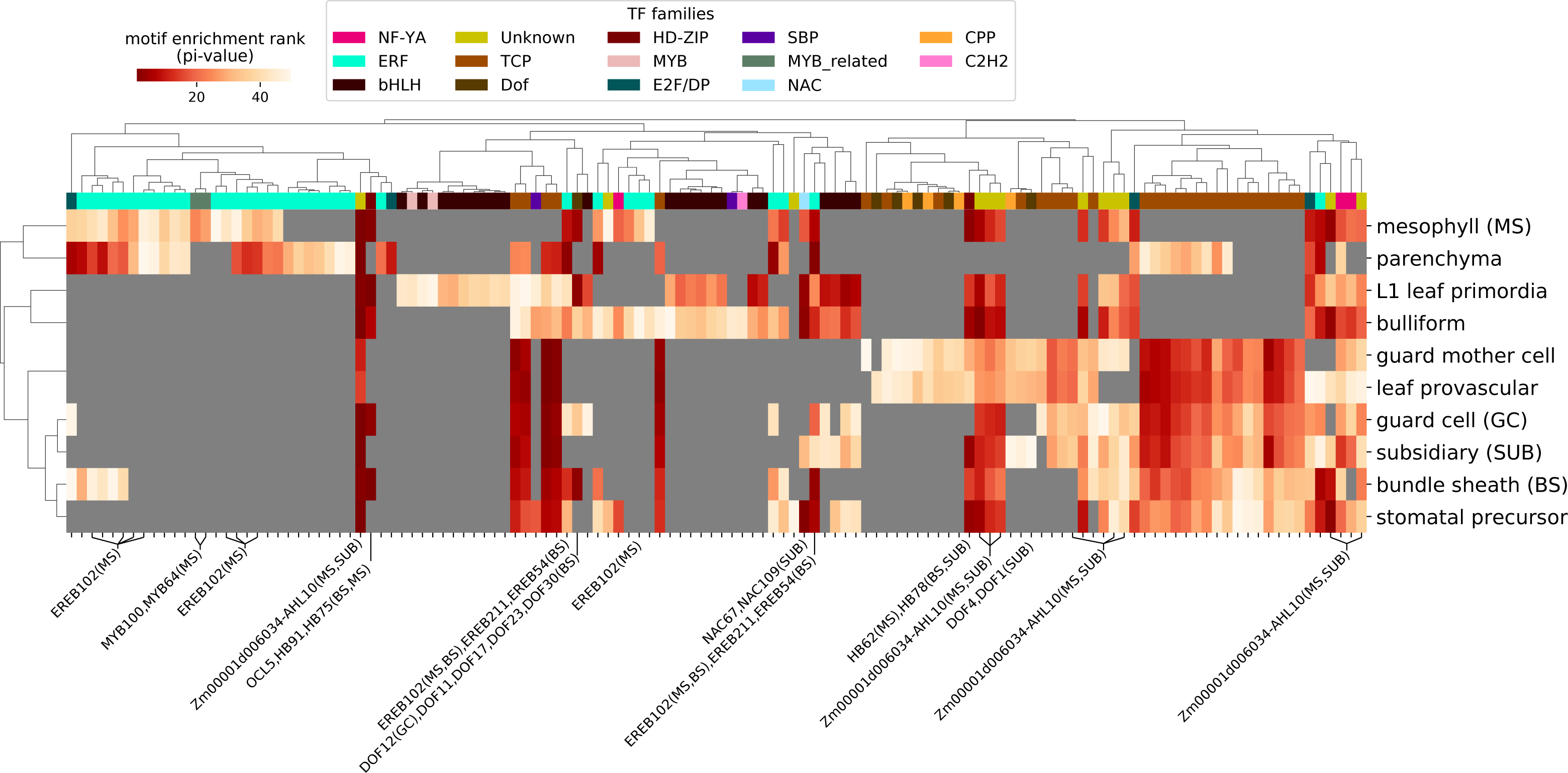
Motif enrichment analysis of a leaf single-cell dataset in maize. Heatmap showing motif enrichment ranks of the top 50 enriched motifs in the leaf cell types of the Marand dataset. On the x-axis, cell-type specific DE TFs associated with the enriched motifs are shown, and, in parentheses, the cell-type where the TF is DE. The column annotation corresponds to different color-coded TF families. Cells in gray indicate motifs enriched in a cell type, but outside the top 50.

For bundle sheath 53 out of 73 (72%) DE TFs were associated with at least one enriched motif. Besides the identification of OCL5a, HB75, and HB91 (Table S7) (Javelle et al. 2010; 2011), three bundle sheath DE TFs ranked third in motif enrichment: DOF11, DOF17, and DOF23. Their Arabidopsis ortholog VDOF1 was characterized in lignin biosynthesis of vascular tissue in Arabidopsis (Ramachandran et al. 2020). In maize, bundle sheath cells are surrounded by a suberin barrier, a molecule that shares precursors with lignin (Mertz and Brutnell 2014; Bezrutczyk et al. 2021). Additionally, DOF motifs have been previously shown to be preferentially enriched in bundle sheath cells (in comparison with mesophyll), further suggesting these are relevant candidates with an important role in bundle sheath function (Dai et al. 2022). Examples of other TFs identified for bundle sheath can be found in Table S7.

Regarding guard cell, we found enrichment at rank 35 for the DE TF DOF12, which is an ortholog of Arabidopsis SCAP1, a regulator of the final stages of guard cell maturation (Negi et al. 2013). Functional analysis of the guard cell GRN (Figure S8A) revealed regulons with functional enrichment related with this cell-type’s function, e.g. “regulation of cellular response to alkaline pH” (Suhita et al. 2004), “regionalization”, “regulation of microtubule polymerization or depolymerization” (Y. Li et al. 2022), “response to water deprivation” (Lawson and Matthews 2020), “cellular response to lipid” (Misra et al. 2015) and “movement of cell or subcellular component” (Y. Li et al. 2022). A maize ortholog of SCAP1, DOF25, was predicted to be involved in cellular response to lipids (Negi et al. 2013). JAM1, JAM2 and ZBF11 are Arabidopsis bHLH TFs involved in response to jasmonate, a hormone known to regulate stomatal movement, and their maize orthologs (bHLH57 of JAM1, bHLH99 of JAM2, and MYC7 and bHLH91 of ZBF1) were predicted by MINI-AC to regulate response to alkaline pH in guard cells. Alkalinization of cytoplasm has been observed to precede jasmonate-induced stomatal closure (Suhita et al. 2004).

In subsidiary cells, NAC109 (ortholog of ATAF1) and NAC67 (ortholog of VNI2) were DE, both with rank 32. NAC TFs, in leaf, are mostly known for their major role in senescence (Kim, Nam, and Lim 2016), and ATAF1 and VNI2 have been reported to be involved in this process (Yang et al. 2011; Garapati et al. 2015), among others (Delessert et al. 2005; Wang et al. 2009; Yamaguchi et al. 2010). While NAC67 was up-regulated and NAC109 was down-regulated, their function in subsidiary cells is currently unknown.

Besides validating the top ranked regulators using cell-type specific DE genes, we also investigated if MINI-AC can identify new regulators that are currently lacking experimental support based on expression. We selected all the regulators among the top 10 ranks for each ACR dataset and compared their ranks in the other datasets (Figure S9). The TFs of the TCP family, despite not being DE in any cell type, had a very low enrichment rank in various datasets which is in agreement with previous findings of TCP genes regulating traits of leaf development (Chai et al. 2017). Other TFs, among them DOF and OCL/HD-ZIP TFs, showed the opposite trend: highly enriched in single-cell derived datasets compared to bulk datasets. One of them, OCL3, has been correlated with leaf length and width (Cruz et al. 2020). We found four motifs from the CPP TF family that showed low ranks in leaf provascular, guard mother cells, and subsidiary cells (Figure 5; the motif with the lowest ranks has 33, 34, and 49 respectively). Different of these CPP motifs are associated with TFs that are orthologs of Arabidopsis TSO1 (as well as SOL1 and SOL2; Figure S8B), which is involved in the stomatal lineage specification (Simmons et al. 2019).

Finally, we explored if MINI-AC was able to capture the known functional partition between mesophyll and bundle sheath cells in maize C4 metabolism (Edwards et al. 2001) using single-cell-derived ACRs. Their respective GRNs were filtered for DE genes in each cell-type and then used for functional enrichment. For both cell types, we identified GO terms related with their specific photosynthetic functions, such as carbon utilization (mesophyll) or RuBisCO subunits assembly (bundle sheath), among others (Figure S10 and Table S8) (Majeran et al. 2005; Dai et al. 2022). Additionally, by selecting regulons showing function enrichment for photosynthesis, we observed that the photosynthetic GRNs of each cell-type have many shared elements (80% of TGs shared; Figure S11A). We also identified putative overlapping regulators of genes known to be cell-type specific for both cell-types, like carbonic anhydrases for mesophyll and RuBisCO subunits for bundle sheath (several NAC and HB TFs, as well as KANADI3, GLK24, and YABBY5; see Figure S11B).

Taken together, we demonstrated that MINI-AC can be used to retrieve known and novel regulators from both single-cell and bulk ACR data. By ranking motifs based on their significance values in different ACR datasets, MINI-AC identified TFs preferentially enriched in specific leaf cell types, which are DE and were predicted to regulate TGs enriched in biological processes specific to those cell types. We also predicted novel TFs that were not previously characterized with a leaf function, and revealed that genes that are key for the C4 metabolism in mesophyll and bundle sheath are potentially regulated by a set of partially overlapping TFs.

## Discussion

Identifying and characterizing regulatory DNA is crucial to understand the control of gene expression underlying developmental and adaptive processes in plants. Methods that profile regulatory DNA, such as ATAC-seq, have been developed and applied to plants, both at the bulk and single-cell level (Lu et al. 2019; Marand et al. 2021). To harness the increasing availability of chromatin accessibility datasets for improving our understanding of gene regulation, we developed MINI-AC, a computational method to infer GRNs in plants by integrating TF motif information with chromatin accessibility.

Several methods that integrate TFBS information with chromatin accessibility have been developed. TOBIAS uses ATAC-seq data to identify TF footprints and integrates TFBS to estimate differential binding scores between conditions. In comparison to MINI-AC, TOBIAS is specific for ATAC-seq data, impeding the analysis of accessibility datasets profiled with other methods. TOBIAS identifies footprint scores for different TFBS, meaning it identifies TFs with clear footprints, but this does not necessarily reflect enrichment of that TF in the ACR set. Besides, the tool it uses to map TFBS within the footprints is MOODs (Motif Occurrence Detection Suite; (Korhonen et al. 2009)), which has a high false positive rate (Kulkarni, Marc Jones, and Vandepoele 2019). MEDEA performs motif enrichment using area under the receiver operating characteristic curve, but it does not perform inference of GRN and lacks support for plants. Other methods integrate TFBS and AC data to constrain expression-based network inference. For example, CellOracle, IReNA, and Inferelator 3.0, use accessibility-based GRNs that are refined or extended using single-cell expression data. These GRNs are built by simply scanning TFBS in ACRs, without performing motif enrichment to filter spurious or false positive motif matches. MINI-AC, conversely, finds the motifs that occur in an ACR set more than expected by chance, and exclusively considers these for the GRN inference.

When comparing the motif enrichment performances of different tools, MINI-AC outperformed Giggle and Bedtools in predicting cell-type specific DE genes given cell-type specific ACRs. MINI-AC proves a better alternative for real accessibility datasets with complex motif signals. This could be explained by differences in the enrichment methodology, given that MINI-AC computes a p-value by contrasting against a background distribution of motif matches within ACRs, while Giggle and Bedtools use a Fisher’s Exact two-tailed test considering overlapping and unique intervals. Although this gives Giggle and Bedtools a computational efficiency advantage, our results show that this comes with a reduction in performance in complex datasets. Inference of an Arabidopsis GRN for mesophyll showed the power of MINI-AC to correctly identify known regulators of circadian rhythm and light perception, demonstrating MINI-AC’s potential to predict novel regulators.

Maize has a genome with long intergenic regions and a high abundance of transposable elements (Jiao et al. 2017), resulting in a complex landscape of regulatory DNA (Lu et al. 2019). Therefore, we evaluated the impact of using different maize non-coding genomic spaces on GRN inference. We observed that locus-based MINI-AC with a small non-coding genomic space yielded the most complete GRNs (0.139 recall and 0.665 precision), while the genome-wide mode predicted the most precise GRNs (0.121 recall and 0.736 precision), with 28% of its ChIP-confirmed interactions being distal. This is important for maize, where approximately one third of the ACRs are distal (Lu et al. 2019). Although it seems counterintuitive that smaller non-coding genomic spaces produce larger GRNs, this is due to a strict FIMO internal cut-off to prevent spurious motif matches when working in the genome-wide mode. Larger non-coding genomic spaces have a lower density of functional TFBS, so FIMO applies more stringent internal cut-offs, yielding less motif matches, especially for low-complexity motifs. Nevertheless, the genome-wide mode prioritizes distal motif matches if they have a higher score than proximal motif matches. Although masking of repetitive sequences and transposable elements during motif mapping could be an alternative to improve genome-wide performance, transposable elements frequently contain binding motifs for transcription factors (Hénaff et al. 2014) and transposons within ACRs have been associated with higher expression of nearby genes (Noshay et al. 2021), indicating their importance for wiring transcriptional networks (Qiu and Köhler 2020).

Using MOA-seq peaks as MINI-AC’s input showed better performance at predicting ChIP-confirmed regulatory interactions compared to using MOA-seq footprints, but the latter showed a better prioritization of relevant motifs at low ranks. When evaluating ChIP-confirmed interactions, the MINI-AC GRN inferred using AC from MOA-seq showed lower precision, but higher recall compared to the GRN inferred using AC from ATAC-seq, which resulted in a MOA-seq F1 between 0.245-0.275 and a ATAC-seq F1 of 0.207. The difference in recall is expected given that the MOA-seq dataset was composed of 593,781 peaks (56 Mbps) and 214,198 footprints (32 Mbps) in comparison to the ATAC-seq dataset with 32,481 peaks (13 Mbps), but the precision of MOA-seq did not drop below 0.5, meaning it can still control for false positives. This proves MOA-seq as a valuable method to learn chromatin accessibility-based GRNs.

We demonstrated that MINI-AC works with single-cell-derived data using the dataset of (Marand et al. 2021) on different leaf cell types. MINI-AC predicts enriched motifs that are associated with cell-type specific DE TFs, and this number increases among the top-ranked motifs. Moreover, a comparison of motif enrichment ranks between bulk and single-cell datasets allowed to identify general and putative cell-type specific regulators of leaf. The application of MINI-AC to study the C4 metabolism in maize revealed that the photosynthetic GRNs of mesophyll and bundle sheath had many shared elements, but they allowed to pinpoint putative regulators of processes and genes specific to each cell-type. On one hand, single-cell ATAC-seq provides accessibility landscapes at single-cell resolution, allowing to find cell-type specific ACRs that can be used to predict cell-type specific GRNs. On the other hand, bulk accessibility profiling still holds value to identify master regulators that do not show a cell-type specific expression pattern. Yet, a direct comparison of the suitability of bulk or single-cell for GRN inference is challenging due to the lack of a gold standard reporting cell-type specific interactions. Additionally, MINI-AC’s evaluation on leaf was done using the availability of a map of mesophyll regulatory interactions for 104 TFs profiled with ChIP-seq (Tu et al. 2020), allowing to benchmark the predicted GRNs. As the profiling of regulatory interactions in maize is extended to different tissues, further evaluations assessing cell-type specific GRNs will be possible.

One of MINI-AC’s limitations is the incomplete characterization of TF motifs, especially for plants (Jaime A Castro-Mondragon et al. 2022). We tried to alleviate this bottleneck in the direct determination of TFBS by using TF motifs predicted with similarity regression of sequence specificity of DNA-binding domains (Weirauch et al. 2014; Lambert et al. 2019). This, however, produces an elevated number of TFs associated to the same motif, and vice versa, which makes the identification of candidate regulators challenging. Nevertheless, given the predictive power of TF motifs for building GRNs (Wilkins et al. 2016; Kulkarni et al. 2018; Kajala et al. 2021; Ferrari, Manosalva Pérez, and Vandepoele 2022), it is crucial that additional efforts are directed towards the direct experimental profiling of TFBS in plants for the yet uncharacterized TFs. In the meanwhile, our results indicate that integrating gene expression information is a good alternative for the prioritization of TF candidates by filtering out TFs not expressed in a specific organ, cell type or condition. In the MINI-AC locus-based mode, the motif matches are automatically assigned to TGs. For the genome-wide mode, the motif matches are annotated to the closest TG, and we showed that annotating the closest gene has a higher ratio of annotation of DE genes vs. non-DE genes, compared to annotating the two closest genes. However, in maize, there are important phenotypic traits controlled by distal regulatory interactions spanning hundreds of kilobases and several intermediate genes (Peng et al. 2019). These are currently not captured by MINI-AC, but the recent application of ChIA-PET (G. Li et al. 2010) to maize (Peng et al. 2019; E. Li et al. 2019), a technology that profiles genome-wide long-range chromatin interactions, could be used to integrate these interactions into the GRN predictions.

With the continuous progress of omics and single-cell technologies, the development of algorithms that effectively exploit this data is crucial (Depuydt, De Rybel, and Vandepoele 2023). Specifically, integrating accessibility and expression data from the same cell-type or cells enables adding new layers of information to the GRNs, such as the interaction type (activation or repression) or finding cell-to-cell mobile TFs (Marand et al. 2021). Several methods use accessibility-based networks as a scaffold to filter based on expression (e.g. CellOracle (Kamimoto et al. 2023)), so MINI-AC is a promising tool that can be used to generate input ACR-based GRNs for such tools. As we recently developed MINI-EX (Motif-Informed Network Inference method based on single-cell EXpression data) (Ferrari, Manosalva Pérez, and Vandepoele 2022), a method that integrates single-cell transcriptomic data with TF motif information to define GRNs, the combination of both tools offers new opportunities to infer more accurate cell-type specific GRNs.

In conclusion, we present MINI-AC as a valuable tool to infer chromatin accessibility- and motif-based GRNs, optimized for plants. MINI-AC can be applied to either bulk- or single-cell-derived datasets, inferring accurate GRNs that contain both proximal and distal regulatory interactions. Functional analysis of the different regulons allowed to pinpoint known and novel putative regulators controlling circadian rhythm and light perception in Arabidopsis, and different photosynthetic processes in maize. With the increasing availability of bulk and single-cell chromatin accessibility datasets, MINI-AC is a valuable tool for unraveling regulatory cascades in plant biology.

## Materials and methods

### Integration and curation of TF motifs

For Arabidopsis, TF motifs modeled as position weight matrices (PWMs) were collected from a previously curated collection (Kulkarni, Marc Jones, and Vandepoele 2019) and combined with motifs from JASPAR 2020 (Fornes et al. 2020) and CisBP version 2.00 (Weirauch et al. 2014). The PWMs were compared in a pairwise manner to remove duplicates using the Regulatory Sequence Analysis Tools (RSAT) (Jaime Abraham Castro-Mondragon et al. 2017) program “compare-matrices”. PWMs with a Ncor (similarity metric) of 1 were considered duplicates. The Arabidopsis gene models of the TFs collected are from the gene annotation version Araport11 (Cheng et al. 2017). For maize, TF motifs were collected only from JASPAR 2020 and CisBP 2.00 and the duplicate removal was done as in Arabidopsis. The maize JASPAR 2020 TF gene names associated with each motif were converted manually to AGPv4 gene ID nomenclature by querying MaizeGDB (Woodhouse et al. 2021). The maize CisBP 2.00 TF gene IDs were converted from AGPv3 to AGPv4 gene ID nomenclature using a gene ID conversion table derived from the combination of GRAMENE (Tello-Ruiz, Jaiswal, and Ware 2022) (http://www.gramene.org; accessed 4th October 2021) and MaizeGDB. For both species, the motif/TF family information was retrieved from PlantTFDB (Jin et al. 2017) and PlnTFDB (Riaño-Pachón et al. 2007). The total number of collected motifs was 1699 and 1335, which are associated with 1117 (representing 57 TF families) and 1234 (representing 41 TF families) TFs for Arabidopsis and maize, respectively.

### Extraction of non-coding genomic search space and motif mapping

The extraction of the non-coding genomic search space for the genome-wide MINI-AC mode was done by taking the genomic complement of the CDS features in the genome annotation files of Arabidopsis and maize. The genome versions used were Araport11 (Cheng et al. 2017) for Arabidopsis and AGPv4 (Jiao et al. 2017) for maize, both downloaded from PLAZA monocots 4.5 (Van Bel et al. 2018). The extraction of the non-coding genomic search space for the locus-based MINI-AC mode was done by obtaining, per locus (only protein-coding genes), the 5 kb upstream of the TrSS, introns, and 1 kb downstream of the TrES. For maize, two additional locus-based definitions were also tested: 1 kb upstream of the TrSS, introns, and 1 kb downstream of the TrES (small non-coding genomic search space), and 15 kb upstream of the TrSS, introns, and 2.5 kb downstream of the TrES (large non-coding genomic search space). In all cases, when the promoter or terminator region of one locus overlapped with the coding sequence of another neighboring locus, the regulatory sequences were shortened to avoid overlap. Thus, many regulatory sequences are shorter than the specified window size. In Arabidopsis, for example, the mean upstream promoter size is 2021 bp, and for maize they are 978 bp, 4326 bp, and 11,053 bp for the small, medium and large non-coding genome definitions, respectively. CB (CB version Compiled on Sep 22, 2017) (Frith, Li, and Weng 2003) and FIMO (version 4.11.4) (Grant, Bailey, and Noble 2011) were used to map the motifs on the non-coding genome. Before the motif mapping with CB the PWMs were scaled to 100. The command lines options used for each tool were “fimo -o $output $PWMfile $seqFile” and “cbust-linux $PWMfile $seqFile -c 0 -f 1”.

### Bulk and single-cell chromatin accessibility datasets

The Arabidopsis phloem and epidermis cell-type specific bulk datasets were downloaded from the supplementary dataset 2 of (Tian et al. 2021) and the induced ACRs for phloem and epidermis were selected. For Arabidopsis, bulk differentially accessible chromatin regions between mesophyll and stem cells were obtained from (Kulkarni, Marc Jones, and Vandepoele 2019) (original study (Sijacic et al. 2018)). The number of peaks for each of the Arabidopsis datasets are summarized in Table S1. The maize leaf bulk ACR datasets used were downloaded from GEO (GSE128434 (Lu et al. 2019)) or PlantCADB (PRJNA391551, samples 284 and 285, and PRJNA518749, samples 288 and 289) (https://bioinfor.nefu.edu.cn/PlantCADB/) (K. Ding et al. 2022). The replicates were processed by keeping peaks that overlap at least 50% of their length in either of the replicates, yielding 20,919 peaks for PRJNA391551 and 51,835 peaks for PRJNA518749. Leaf cell-type specific peaks were obtained from a single-cell ATAC-seq maize dataset (Marand et al. 2021). The count per million (CPM) matrix was downloaded from GEO (GSE155178). This matrix contains the CPM value of 165,913 peaks (rows) in 92 different subclusters (columns) grouped in 10 clusters, and annotated to 56 different cell types. The peak specificity per subcluster was estimated by computing the z-score per row (accessibility of peak X in cell type Y - mean accessibility of peak X in all cell types / standard deviation of accessibility of peak X in all cell types). To ensure completeness of the GRNs, we selected the top 10,000 z-scoring peaks of 18 leaf-related subclusters that had > 50% of seedling cells (it was the only sample profiled in (Marand et al. 2021) that contained leaf tissue). These 18 subclusters span 4 clusters and 10 cell type annotations (bundle sheath, mesophyll, bulliform, leaf provascular, subsidiary, guard mother cell, guard cell, stomatal precursor, L1 leaf primordia and parenchyma).

### MOA-seq data processing and peak calling

To assess putative TF footprints genome-wide, we performed MOA-seq on B73 plants grown under long-day conditions (16h day/8h night, 28/21L) for 26 days. For each biological replicate, leaf blade tissue of 12 plants was harvested, pooled and immediately frozen in liquid nitrogen. Collected tissues were homogenized and processed for MOA-seq library constructions following a previous protocol (Savadel et al. 2021).

For the MOA-seq data analysis, the B73 AGPv5 genome (Hufford et al. 2021) was indexed using STAR (v2.7.9a) (Dobin et al. 2013). Raw reads of MOA-seq data were preprocessed using SeqPurge (v2022-07-15) (Sturm, Schroeder, and Bauer 2016) with parameters of “-min_len -qcut 0” Overlapping MOA-seq paired-end reads were merged into single-end reads, including base quality score correction, using NGmerge (v0.3) (Gaspar 2018) with parameters “-p 0.2 -m 15 -d -e 30 -z -v”. Processed reads were aligned against the indexed reference genome using STAR (v2.7.9a) with BAM format output. As STAR is designed to map RNA, we set the flag --alignIntronMax 1 for DNA (prevents introns being allowed). Alignment fragments with less than 81 bp and a MAPQ of 255 (uniquely mapping) were retained for further analysis.

For MOA peak calling, we first merged the replicate’s BAM outputs, determine the average fragment length using samtools (v1.16) (H. Li et al. 2009), and the effective genome size with unique-kmers.py (https://github.com/dib-lab/khmer/). MACS3 (v3.0.0a7) (Zhang et al. 2008) was used to determine significant peaks with parameters “-q 0.05 -g {effective genome size} -s {fragment length} --min-length {fragment length} -max-gap {2x fragment length} --nomodel --extsize {fragment length} --keep-dup all”. To obtain high resolution MOA footprints, reads were shortened to 20bp around the center of each read as described in (Liang et al. 2022) and significant footprints were determined with MACS3 using the same parameters as were used for the peak files. For comparison with AGPv4 data, MACS3 peaks were converted to B73 AGPv4 using liftOver (Hinrichs et al. 2006), with default parameters, using the chain file from Ensembl (Cunningham et al. 2022). The MOA-seq raw sequencing data has been deposited to the NCBI under the BioProject ID PRJNA961163 (BioSamples IDs SAMN34353189, SAMN34353190, and SAMN34353191).

### Motif enrichment in ACRs and inferring TF - target gene regulatory interactions

To estimate the motif matches in ACR expected by chance, the input ACRs are shuffled 1000 times across the non-coding genomic search space and the number of motif matches overlapping with them are counted. The procedure of shuffling peaks across the non-coding regions did not show differences when considering the GC percentage of the peaks in the ACR background generation (4 bins of the non-coding genome regions based on their GC percentage were generated and each peak of the input ACR set was shuffled only across the corresponding bin). Therefore, we kept the shuffling across all the non-coding regions at once. An empirical p-value is computed by counting how many times the number of motif matches in ACRs expected by chance is equal or larger than the real one. The enrichment fold is computed as the real number of motif matches in ACR divided by the number of motif matches in ACR expected by chance. The p-value is corrected for multiple hypothesis testing using the Benjamini-hochberg method (q-value) (Benjamini and Hochberg 1995)). TFs associated with enriched motifs (q-value below threshold) are linked to their target genes if a TFBS within an ACR peak is in the regulatory sequence of a gene (locus-based mode) or in its proximity (genome-wide mode). To rank the motifs, the π-value was used, which is the -log10(q-value) multiplied by the enrichment fold.

In the locus-based MINI-AC mode, the TFs are linked to their corresponding TGs when there is a motif match within an ACR. In the genome-wide MINI-AC mode, the motif matches within ACRs are annotated to the closest gene using “bedtools closest –a $motif_matches_within_ACR -b $genes_coordinates -t all -d -D b -k 1” (Quinlan and Hall 2010). The number of closest genes to be annotated as TGs can be changed to two, in case selecting the closest gene is deemed too restrictive (https://github.com/VIB-PSB/MINI-AC). Additionally, it is possible to set a cut-off distance of the second-closest gene to the peak for it to be annotated as TG to avoid the annotation of very distal second-closest genes that might increase the number of false positives.

The suitability of annotating the second-closest genes as TGs was done by using two maize cell-type specific scATAC-seq datasets (mesophyll and bundle sheath; methods section “Bulk and single-cell chromatin accessibility datasets”) (Marand et al. 2021) and assessing the proportion of first and second-closest genes that were DE. Precision was computed as the number of annotated genes (first only, or first and second) that are DE, divided by the total number of annotated TGs. Recall was computed as the number of DE TGs divided by the total number of DE genes. Precision, recall, and F1 were computed using the ACR peaks annotating only the first closest gene, and then adding to these genes the second-closest genes if they fall within a certain cut-off distance from the gene.

MINI-AC has been implemented as a Nextflow pipeline (https://www.nextflow.io/) (Di Tommaso et al. 2017) version 21.10.6 (DLS2) and can be downloaded from the GitHub repository https://github.com/VIB-PSB/MINI-AC and executed as a command line pipeline following the instructions in the tutorial. Examples of the different motif enrichment and network output files are described in a tutorial on the MINI-AC GitHub page.

### GO enrichment of network regulons

For Arabidopsis, the GO annotations were downloaded from http://geneontology.org/gene-associations/tair.gaf.gz on 22nd July 2021. They were filtered for GO annotations of type “full”, and evidence types “experimental” and “author/curator statement” (evidence codes: EXP, IMP, IDA, IPI, IGI, IEP, TAS, NAS, and IC). For maize, the GO annotations were downloaded from PLAZA Dicots 4.5 (https://bioinformatics.psb.ugent.be/plaza/versions/plaza_v4_5_dicots/) (Van Bel et al. 2018). GO annotation files were extended for parental terms and filtered for GO terms that were associated with less than 1000 genes (discarding very general GO terms), and for GO terms of the category “Biological Process” (BP). The GO enrichment of the networks predicted by MINI-AC is calculated using the hypergeometric distribution to determine a q-value by correcting for multiple hypotheses testing with the Benajmini-Hochberg procedure. This means that for each regulon in the network, the overrepresentation of specific biological processes in the TGs is evaluated.

### Comparison of genomic interval enrichment methods in Arabidopsis

To compare MINI-AC with Giggle (Layer et al. 2018) and Bedtools Fisher (Quinlan and Hall 2010), both programs were run with the following parameter setting: “giggle index -i input_file –o giggle_index –f; giggle search -I giggle_index –a acr_file –s –g 119667750” and “bedtools fisher –a motif_mappings –b acr_file –g fasta_index”. The “synthetic” ACR dataset was obtained by intersecting an Arabidopsis seedling ACR dataset (Lu et al. 2017) (downloaded from GEO with accession number GSE85203; 50k and 500 nuclei samples were combined after processing replicates by keeping peaks overlapping reciprocally 50% of their length) with the 500 top scoring ChIP-seq peaks of 19 out 21 TFs profiled in Arabidopsis (Song et al. 2016) for which there was a motif available in our motif collection (downloaded from GEO with accession number GSE80564; only the EtOH samples were used). For each method, the p-values returned were corrected for multiple testing using a Benjamini-Hochberg procedure, yielding q-values. The minimum p-value that MINI-AC can generate for motif enrichment is 0.001 (as a result of obtaining the background ACR by shuffling the real ACR 1000 times), while Giggle and Bedtools Fisher give p-values in very small orders of magnitude (up to 10^-200^). Therefore, we used a set of 100 shuffled datasets (the 3 real ACR and the “synthetic” ACR datasets were shuffled 25 times each using “bedtools shuffle”) where we expected zero enriched motifs (any enriched motif at a specific q-value can be considered a false positive) and determined the q-value at which we obtained zero enriched motifs to use as the q-value cut-off. Using the q-values at an FDR of 0% (0.01 for locus-based MINI-AC, 0.1 for genome-wide MINI-AC, 10^-32^ for Giggle, and 10^-31^ for Bedtools Fisher), precision, recall, and F1 were computed for each method and dataset. Precision was computed as the number of enriched motifs belonging to true positive TFs divided by the total number of enriched motifs predicted. Recall was computed as the number of enriched motifs belonging to true positive TFs divided by the total number of motifs belonging to true positive TFs. F1 was computed as the harmonic mean between precision and recall.

### Comparison of MINI-AC non-coding genomic search space strategies in maize

The motif enrichment results for the different MINI-AC modes and non-coding genomic search space definitions were evaluated by ranking motifs using the π-value. The resulting networks were evaluated, by computing precision, recall, and F1, using a previously published maize leaf gold standard of TFBS of 62 TFs profiled using ChIP-seq (Tu et al. 2020). The ChIP-seq peaks were annotated to the closest gene using “bedtools closest” to obtain a ChIP-seq-based GRN. To benchmark each MINI-AC non-coding space definition and mode with the possibility of obtaining a recall of 1, the gold standard was adapted accordingly (so-called “adapted GS”). The gold standard network was first filtered for TFs associated with motifs present in our PWM collection. Then, the peaks that did not contain the motif of the correct incoming TF were removed, based on FIMO and CB motif mapping. To determine the proportion of correct TF-TG interactions predicted by each MINI-AC mode and non-coding genomic search space definition, the motif mappings within ACRs were overlapped with the “adapted GS” peaks, and its distance to the corresponding target gene was determined using “bedtools closest”.

## Acknowledgements

We would like to thank Sander Thierens and Bert Droesbeke for their assistance building the Docker image, Jasper Staut for testing the pipeline, Svitlana Lukicheva for proofreading the GitHub repository tutorial, and Dries Vaneechoutte for his work on compiling the hypergeometric enricher used.

## Accession numbers

The raw and processed MOA-seq data used in this work has been deposited to the NCBI under the BioProject ID PRJNA961163 (BioSamples IDs SAMN34353189, SAMN34353190, and SAMN34353191).

## Data availability statement

All relevant data can be found within the article and its supporting information.

## Author contributions

K.V. and N.M.P. conceived the project and designed the research. C.F and N.M.P. collected and curated the PWM collection. K.V. and N.M.P developed the method and performed the analyses. J.E. and T.H. generated and processed the MOA-seq data. T.D and H.N proofread, edited and provided input on the manuscript. N.M.P. and K.V. wrote the manuscript. All authors read and approved the final paper.

## Supplementary Figure 1

**Supplementary Figure 1. Genome coverage of Arabidopsis and maize using the non-coding genomic space extraction strategies of locus-based MINI-AC mode.** Pie charts showing the percentage of non-coding genome covered by the extraction strategy of the MINI-AC locus-based mode in Arabidopsis (A) and maize (B) using 5 kb upstream of the TrSS, introns, and 1 kb downstream of the TrES. Black represents the part of the non-coding genome that is covered, and white represents the part that is not covered. In parentheses, there is the absolute number of mega base-pairs represented by each percentage.

## Supplementary Figure 2

**Supplementary Figure 2. Calibration of optimal top-scoring motif matches from CB and FIMO in Arabidopsis and maize.** (A) Proportion of TGs targeted by each motif on FIMO-derived motif mappings, using all matches, top 7,000 matches and top 16,000 matches in Arabidopsis and/or maize. (B) Performance statistics precision, recall, and F1 for different top-scoring sets of motif matches for FIMO and CB in Arabidopsis and maize by overlapping them with a ChIP-seq derived gold standard set of TFBS.

## Supplementary Figure 3

**Supplementary Figure 3. Assessment of second-closest gene annotation strategy.** Performance statistics for retrieval of cell-type specific DE genes –(A) mesophyll, (B) bundle sheath–, when annotating only the closest gene to a peak and adding progressively the second-closest genes that fall within a certain distance cutoff (x-axis).

## Supplementary Figure 4

**Supplementary Figure 4. Benchmark of locus-based and genome-wide MINI-AC modes on maize.** Area under the precision-recall curve (AUPRC) for the retrieval of enriched motifs by MINI-AC that are associated with TFs of the maize leaf gold standard, for the four MINI-AC modes and non-coding genomic search spaces tested (locus-based or genome-wide).

## Supplementary Figure 5

**Supplementary Figure 5. Evaluation and comparison of motif enrichment and GRNs predicted by MINI-AC using peaks and footprints derived from maize leaf MOA-seq.**

(A) Precision-recall curve obtained by evaluating ranked motif enrichment results in MOA-seq peaks and footprints using a set of combined DE cell-types specific genes. (B) AUPRC relative to precision-recall curve of panel A. (C) Venn diagram showing the number of unique and shared enriched motifs predicted from MOA-seq peaks and footprints. (D) Venn diagram of the number of unique and shared TF-TG interactions of GRNs predicted using MOA-seq peaks and footprints. (E) Venn diagram of the number of unique and shared ChIP-confirmed TF-TG interactions of GRNs predicted using MOA-seq peaks and footprints.

## Supplementary Figure 6

**Supplementary Figure 6. Comparison of ATAC-seq peaks derived from bulk and single-cell technologies.** (A) Distribution plot of the distance to the closest gene for the indicated bulk ACR datasets. (B) Distribution plot of the distance to the closest gene for all the peaks of the Marand dataset. (C) Distribution plot of the distance to the closest gene for the top 10,000 z-scoring peaks of the Marand dataset. (D) Bar plot showing the median z-score (proxy for cell-type specificity) for the top 10,000 z-scoring peaks of the Marand dataset, previously binned by distance to the closest gene.

## Supplementary Figure 7

**Supplementary Figure 7. UpSet plot comparing the total motifs and motifs associated with DE TFs predicted for different bulk and single-cell leaf datasets.** (A) Comparison of the shared and unique enriched motifs for the different bulk and single-cell ACR sets. For visualization purposes, we only show overlapping sets with more than five elements. (B) Comparison of the shared and unique enriched motifs associated with DE TFs for the different bulk and single-cell ACR sets. For visualization purposes, we only show overlapping sets with more than 5 elements.

## Supplementary Figure 8

**Supplementary Figure 8. Functional network analysis of guard cells and rank distribution of TSO1 ortholog in maize**. (A) Heatmap representing the GO enrichment of the MINI-AC GRN for guard cells (all genes, DE and not DE). The GO term “cellular response to lipid” enriched for several DOF TFs, but for visualization purposes only DOF25, a maize ortholog of the Arabidopsis gene SCAP1, was kept, as SCAP1 is a known regulator of Arabidopsis guard cell function. (B) Bar plot showing the rank distribution of motifs associated with TSO1, SOL1 and SOL2 on different leaf cell types of maize.

## Supplementary Figure 9

**Supplementary Figure 9. Comparison of motif enrichment ranks across different bulk and single-cell ACR sets of maize leaf.** Bubble plot showing the rank of all the TFs associated with motifs within the top 10 enrichment rank of any dataset in all the different bulk and single-cell ACR sets. The rows are colored based on the DE status of the TF on different leaf cell types, while the bubble size represents the motif enrichment rank. The lower the rank, the bigger the bubble.

## Supplementary Figure 10

**Supplementary Figure 10. Functional analysis of maize mesophyll and bundle sheath GRNs predicted by MINI-AC.** Heatmap summarizing the GO enrichment significance (measured by -log10(q-value) in orange) of bundle sheath and mesophyll MINI-AC regulons GRNs, considering only DE TF and TGs. The row annotations are, in blue, the number of TFs with such enriched GO terms (mean if the enrichment is in the two cell types) and, in purple, the number of TGs annotated with this GO term regulated by the TF (mean if the enrichment is in the two cell types).

## Supplementary Figure 11

**Supplementary Figure 11. Photosynthesis GRNs of mesophyll and bundle sheath predicted by MINI-AC.** The node color indicates in which cell types the genes are DE: green for mesophyll, orange for bundle sheath, and purple for both. The edge color indicates in which GRN the interaction is present: green for mesophyll, orange for bundle sheath, and purple for both. The diamond-shaped nodes represent regulators, while the circle-shaped nodes represent target genes. (A) Mesophyll and bundle sheath GRNs predicted by MINI-AC, filtered for TFs that are DE in those specific cell types and TGs that are annotated to “photosynthesis” GO term or child terms of it. The nodes have been distributed, so the bundle sheath DE TFs are in the upper right part, the mesophyll TFs are in the upper left part, and the DE TFs in both cell types are down. The TGs controlled by the three groups of TFs are in the center. The TFs controlled uniquely by one group of TFs are in the outer part. The TGs shared between two groups of TFs are found in between the corresponding TF groups. (B) Regulators predicted by MINI-AC for carbonic anhydrase 1, 2, and 6, and RuBisCO subunits 1 and 2.

## Supplementary Table 1

**Supplementary Table 1. Summary of datasets used for MINI-AC benchmark in Arabidopsis.**

## Supplementary Table 2

**Supplementary Table 2. Statistics of the TF-TG leaf-specific gold standard set adapted to each MINI-AC mode and non-coding genomic space definition.**

## Supplementary Table 3

**Supplementary Table 3. Summary of the motif enrichment results and GRNs predicted by MINI-AC using two types of ACRs derived from MOA-seq.**

## Supplementary Table 4

**Supplementary Table 4. Summary of the motif enrichment results for leaf bulk- and single-cell-derived ACR sets.**

## Supplementary Table 5

**Supplementary Table 5. AGPv4 gene IDs of the genes mentioned in the publication.**

## Supplementary Table 6

**Supplementary table 6. Summary of the motif enrichment ranks for cell-type specific DE TFs in different bulk- and single-cell-derived ACR sets, along with metadata and expression data about those TFs.**

## Supplementary Table 7

**Supplementary Table 7. Summary of examples of regulators and regulons predicted by MINI-AC which show literature support.**

## Supplementary Table 8

**Supplementary Table 8. MINI-AC-predicted regulators with regulons showing enrichment of GO terms related with C4 metabolism.**

## Supplementary Dataset 1

**Supplementary Dataset 1. MINI-AC TF-centered table, motif-centered table, functional network and node attributes table result files for the Arabidopsis dataset of mesophyll used in this study (Sijacic et al. 2018).**

## Supplementary Dataset 2

**Supplementary Dataset 2. GRN predicted from cell-type specific peaks of the Marand dataset, reporting cell-type specific interactions in mesophyll, bundle sheath, guard cells and subsidiary cells. It includes functional enrichment information.**

## Supplementary Dataset 3

**Supplementary Dataset 3. MINI-AC TF-centered table and motif-centered table result files for the Marand dataset of the cell types mesophyll, bundle sheath, subsidiary and guard cell.**

## Supplementary Methods 1

**Supplementary Methods 1. Calibration of the optimal set of top-scoring motif mappings**

## Supplementary Methods 2

**Supplementary Methods 2. Cell-type specific DE genes**

## Supplementary Methods 3

**Supplementary Methods 3. Leaf-expressed genes in maize**

## Literature cited

Aibar, Sara, Carmen Bravo González-Blas, Thomas Moerman, Vân Anh Huynh-Thu, Hana Imrichova, Gert Hulselmans, Florian Rambow, et al. 2017. “SCENIC: Single-Cell Regulatory Network Inference and Clustering.” Nature Methods 14 (11): 1083–86. https://doi.org/10.1038/nmeth.4463.

Angerer, Philipp, Lukas Simon, Sophie Tritschler, F. Alexander Wolf, David Fischer, and Fabian J. Theis. 2017. “Single Cells Make Big Data: New Challenges and Opportunities in Transcriptomics.” Current Opinion in Systems Biology, Big data acquisition and analysis • Pharmacology and drug discovery, 4 (August): 85–91. https://doi.org/10.1016/j.coisb.2017.07.004.

Banf, Michael, and Seung Y. Rhee. 2017. “Computational Inference of Gene Regulatory Networks: Approaches, Limitations and Opportunities.” Biochimica et Biophysica Acta - Gene Regulatory Mechanisms 1860 (1): 41–52. https://doi.org/10.1016/j.bbagrm.2016.09.003.

Benjamini, Yoav, and Yosef Hochberg. 1995. “Controlling the False Discovery Rate: A Practical and Powerful Approach to Multiple Testing.” Journal of the Royal Statistical Society. Series B (Methodological) 57 (1): 289–300.

Bentsen, Mette, Philipp Goymann, Hendrick Schultheis, Kathrin Klee, Anastasiia Petrova, René Wiegandt, Annika Fust, et al. 2020. “ATAC-Seq Footprinting Unravels Kinetics of Transcription Factor Binding during Zygotic Genome Activation.” Nature Communications, no. 2020. https://doi.org/10.1038/s41576-018-0087-x.

Bezrutczyk, Margaret, Nora R. Zöllner, Colin P.S. Kruse, Thomas Hartwig, Tobias Lautwein, Karl Köhrer, Wolf B. Frommer, and Ji Yun Kim. 2021. “Evidence for Phloem Loading via the Abaxial Bundle Sheath Cells in Maize Leaves.” Plant Cell 33 (3): 531–47. https://doi.org/10.1093/plcell/koaa055.

Bou-Torrent, Jordi, Mercè Salla-Martret, Ronny Brandt, Thomas Musielak, Jean-Christophe Palauqui, Jaime F. Martínez-García, and Stephan Wenkel. 2012. “ATHB4 and HAT3, Two Class II HD-ZIP Transcription Factors, Control Leaf Development in Arabidopsis.” Plant Signaling & Behavior 7 (11): 1382–87. https://doi.org/10.4161/psb.21824.

Boyle, Alan P., Sean Davis, Hennady P. Shulha, Paul Meltzer, Elliott H. Margulies, Zhiping Weng, Terrence S. Furey, and Gregory E. Crawford. 2008. “High-Resolution Mapping and Characterization of Open Chromatin across the Genome.” Cell 132 (2): 311–22. https://doi.org/10.1016/j.cell.2007.12.014.

Buenrostro, Jason D., Beijing Wu, Howard Y. Chang, and William J. Greenleaf. 2015. “ATAC-Seq: A Method for Assaying Chromatin Accessibility Genome-Wide.” Current Protocols in Molecular Biology 109 (January): 21.29.1-21.29.9. https://doi.org/10.1002/0471142727.mb2129s109.

Castro-Mondragon, Jaime A, Rafael Riudavets-Puig, Ieva Rauluseviciute, Roza Berhanu Lemma, Laura Turchi, Romain Blanc-Mathieu, Jeremy Lucas, et al. 2022. “JASPAR 2022: The 9th Release of the Open-Access Database of Transcription Factor Binding Profiles.” Nucleic Acids Research 50 (D1): D165–73. https://doi.org/10.1093/nar/gkab1113.

Castro-Mondragon, Jaime Abraham, Sébastien Jaeger, Denis Thieffry, Morgane Thomas-Chollier, and Jacques Van Helden. 2017. “RSAT Matrix-Clustering: Dynamic Exploration and Redundancy Reduction of Transcription Factor Binding Motif Collections.” Nucleic Acids Research 45 (13): 1–13. https://doi.org/10.1093/nar/gkx314.

Chai, Wenbo, Pengfei Jiang, Guoyu Huang, Haiyang Jiang, and Xiaoyu Li. 2017. “Identification and Expression Profiling Analysis of TCP Family Genes Involved in Growth and Development in Maize.” Physiology and Molecular Biology of Plants 23 (4): 779–91. https://doi.org/10.1007/s12298-017-0476-1.

Chen, Dijun, Wenhao Yan, Liang Yu Fu, and Kerstin Kaufmann. 2018. “Architecture of Gene Regulatory Networks Controlling Flower Development in Arabidopsis Thaliana.” Nature Communications 9 (1): 1–13. https://doi.org/10.1038/s41467-018-06772-3.

Cheng, Chia-Yi, Vivek Krishnakumar, Agnes P. Chan, Françoise Thibaud-Nissen, Seth Schobel, and Christopher D. Town. 2017. “Araport11: A Complete Reannotation of the Arabidopsis Thaliana Reference Genome.” The Plant Journal: For Cell and Molecular Biology 89 (4): 789–804. https://doi.org/10.1111/tpj.13415.

Creux, Nicky, and Stacey Harmer. 2019. “Circadian Rhythms in Plants.” Cold Spring Harbor Perspectives in Biology 11 (9): a034611. https://doi.org/10.1101/cshperspect.a034611.

Cruz, Daniel Felipe, Sam De Meyer, Joke Ampe, Heike Sprenger, Dorota Herman, Tom Van Hautegem, Jolien De Block, Dirk Inzé, Hilde Nelissen, and Steven Maere. 2020. “Using Single-Plant-Omics in the Field to Link Maize Genes to Functions and Phenotypes.” Molecular Systems Biology 16 (12): e9667. https://doi.org/10.15252/msb.20209667.

Cunningham, Fiona, James E Allen, Jamie Allen, Jorge Alvarez-Jarreta, M Ridwan Amode, Irina M Armean, Olanrewaju Austine-Orimoloye, et al. 2022. “Ensembl 2022.” Nucleic Acids Research 50 (D1): D988–95. https://doi.org/10.1093/nar/gkab1049.

Dai, Xiuru, Xiaoyu Tu, Baijuan Du, Pengfei Dong, Shilei Sun, Xianglan Wang, Jing Sun, et al. 2022. “Chromatin and Regulatory Differentiation between Bundle Sheath and Mesophyll Cells in Maize.” The Plant Journal 109 (3): 675–92. https://doi.org/10.1111/tpj.15586.

De Clercq, Inge, Jan Van de Velde, Xiaopeng Luo, Li Liu, Veronique Storme, Michiel Van Bel, Robin Pottie, Dries Vaneechoutte, Frank Van Breusegem, and Klaas Vandepoele. 2021. “Integrative Inference of Transcriptional Networks in Arabidopsis Yields Novel ROS Signalling Regulators.” Nature Plants 2021 7:4 7 (4): 500–513. https://doi.org/10.1038/s41477-021-00894-1.

Delessert, Christian, Kemal Kazan, Iain W. Wilson, Dominique Van Der Straeten, John Manners, Elizabeth S. Dennis, and Rudy Dolferus. 2005. “The Transcription Factor ATAF2 Represses the Expression of Pathogenesis-Related Genes in Arabidopsis.” The Plant Journal 43 (5): 745–57. https://doi.org/10.1111/j.1365-313X.2005.02488.x.

Depuydt, Thomas, Bert De Rybel, and Klaas Vandepoele. 2023. “Charting Plant Gene Functions in the Multi-Omics and Single-Cell Era.” Trends in Plant Science 28 (3): 283–96. https://doi.org/10.1016/j.tplants.2022.09.008.

Di Tommaso, Paolo, Maria Chatzou, Evan W. Floden, Pablo Prieto Barja, Emilio Palumbo, and Cedric Notredame. 2017. “Nextflow Enables Reproducible Computational Workflows.” Nature Biotechnology 35 (4): 316–19. https://doi.org/10.1038/nbt.3820.

Ding, Ke, Shanwen Sun, Yang Luo, Chaoyue Long, Jingwen Zhai, Yixiao Zhai, and Guohua Wang. 2022. “PlantCADB: A Comprehensive Plant Chromatin Accessibility Database.” Genomics, Proteomics & Bioinformatics, October, S1672–0229(22)00133-4. https://doi.org/10.1016/j.gpb.2022.10.005.

Ding, Pingtao, Toshiyuki Sakai, Ram Krishna Shrestha, Nicolas Manosalva Perez, Wenbin Guo, Bruno Pok Man Ngou, Shengbo He, et al. 2021. “Chromatin Accessibility Landscapes Activated by Cell-Surface and Intracellular Immune Receptors.” Journal of Experimental Botany, August. https://doi.org/10.1093/jxb/erab373.

Dobin, Alexander, Carrie A. Davis, Felix Schlesinger, Jorg Drenkow, Chris Zaleski, Sonali Jha, Philippe Batut, Mark Chaisson, and Thomas R. Gingeras. 2013. “STAR: Ultrafast Universal RNA-Seq Aligner.” Bioinformatics 29 (1): 15–21. https://doi.org/10.1093/bioinformatics/bts635.

Dong, Pengfei, Xiaoyu Tu, Po-Yu Chu, Peitao Lü, Ning Zhu, Donald Grierson, Baijuan Du, Pinghua Li, and Silin Zhong. 2017. “3D Chromatin Architecture of Large Plant Genomes Determined by Local A/B Compartments.” Molecular Plant 10 (12): 1497– 1509. https://doi.org/10.1016/j.molp.2017.11.005.

Edwards, Gerald E., Vincent R. Franceschi, Maurice S. B. Ku, Elena V. Voznesenskaya, Vladimir I. Pyankov, and Carlos S. Andreo. 2001. “Compartmentation of Photosynthesis in Cells and Tissues of C 4 Plants.” Journal of Experimental Botany 52 (356): 577–90. https://doi.org/10.1093/jxb/52.356.577.

Ferrari, Camilla, Nicolás Manosalva Pérez, and Klaas Vandepoele. 2022. “MINI-EX: Integrative Inference of Single-Cell Gene Regulatory Networks in Plants.” Molecular Plant 15 (11): 1807–24. https://doi.org/10.1016/j.molp.2022.10.016.

Fornes, Oriol, Jaime A. Castro-Mondragon, Aziz Khan, Robin Van Der Lee, Xi Zhang, Phillip A. Richmond, Bhavi P. Modi, et al. 2020. “JASPAR 2020: Update of the Open-Access Database of Transcription Factor Binding Profiles.” Nucleic Acids Research 48 (D1): D87–92. https://doi.org/10.1093/nar/gkz1001.

Franco-Zorrilla, José M., Irene López-Vidriero, José L. Carrasco, Marta Godoy, Pablo Vera, and Roberto Solano. 2014. “DNA-Binding Specificities of Plant Transcription Factors and Their Potential to Define Target Genes.” Proceedings of the National Academy of Sciences of the United States of America 111 (6): 2367–72. https://doi.org/10.1073/pnas.1316278111.

Frith, Martin C., Michael C. Li, and Zhiping Weng. 2003. “Cluster-Buster: Finding Dense Clusters of Motifs in DNA Sequences.” Nucleic Acids Research 31 (13): 3666–68. https://doi.org/10.1093/nar/gkg540.

Fu, Limin, and Enzo Medico. 2007. “FLAME, a Novel Fuzzy Clustering Method for the Analysis of DNA Microarray Data.” BMC Bioinformatics 8 (1): 3. https://doi.org/10.1186/1471-2105-8-3.

Garapati, Prashanth, Gang-Ping Xue, Sergi Munné-Bosch, and Salma Balazadeh. 2015. “Transcription Factor ATAF1 in Arabidopsis Promotes Senescence by Direct Regulation of Key Chloroplast Maintenance and Senescence Transcriptional Cascades.” Plant Physiology 168 (3): 1122–39. https://doi.org/10.1104/pp.15.00567.

Gardner, Timothy S., and Jeremiah J. Faith. 2005. “Reverse-Engineering Transcription Control Networks.” Physics of Life Reviews 2 (1): 65–88. https://doi.org/10.1016/j.plrev.2005.01.001.

Gaspar, John M. 2018. “NGmerge: Merging Paired-End Reads via Novel Empirically-Derived Models of Sequencing Errors.” BMC Bioinformatics 19 (1): 536. https://doi.org/10.1186/s12859-018-2579-2.

Gaudinier, Allison, and Siobhan M. Brady. 2016. “Mapping Transcriptional Networks in Plants: Data-Driven Discovery of Novel Biological Mechanisms.” Annual Review of Plant Biology 67 (1): 575–94. https://doi.org/10.1146/annurev-arplant-043015-112205.

Gaudinier, Allison, Joel Rodriguez-Medina, Lifang Zhang, Andrew Olson, Christophe Liseron-Monfils, Anne Maarit Bågman, Jessica Foret, et al. 2018. “Transcriptional Regulation of Nitrogen-Associated Metabolism and Growth.” Nature 563 (7730): 259–64. https://doi.org/10.1038/s41586-018-0656-3.

Gibbs, Claudia Skok, Christopher A Jackson, Giuseppe-Antonio Saldi, Aashna Shah, Andreas Tjärnberg, Aaron Watters, Nicholas De Veaux, et al. 2021. “Single-Cell Gene Regulatory Network Inference at Scale: The Inferelator 3.0.” BioRxiv, May, 2021.05.03.442499. https://doi.org/10.1101/2021.05.03.442499.

Grant, Charles E., Timothy L. Bailey, and William Stafford Noble. 2011. “FIMO: Scanning for Occurrences of a given Motif.” Bioinformatics 27 (7): 1017–18. https://doi.org/10.1093/bioinformatics/btr064.

Gray, Jennifer A., Akiva Shalit-Kaneh, Dalena Nhu Chu, Polly Yingshan Hsu, and Stacey L. Harmer. 2017. “The REVEILLE Clock Genes Inhibit Growth of Juvenile and Adult Plants by Control of Cell Size.” Plant Physiology 173 (4): 2308–22. https://doi.org/10.1104/pp.17.00109.

Haque, Samiul, Jabeen S. Ahmad, Natalie M. Clark, Cranos M. Williams, and Rosangela Sozzani. 2019. “Computational Prediction of Gene Regulatory Networks in Plant Growth and Development.” Current Opinion in Plant Biology 47: 96–105. https://doi.org/10.1016/j.pbi.2018.10.005.

Hénaff, Elizabeth, Cristina Vives, Bénédicte Desvoyes, Ankita Chaurasia, Jordi Payet, Crisanto Gutierrez, and Josep M. Casacuberta. 2014. “Extensive Amplification of the E2F Transcription Factor Binding Sites by Transposons during Evolution of Brassica Species.” The Plant Journal: For Cell and Molecular Biology 77 (6): 852–62. https://doi.org/10.1111/tpj.12434.

Heyndrickx, Ken S., Jan Van de Velde, Congmao Wang, Detlef Weigel, and Klaas Vandepoele. 2014. “A Functional and Evolutionary Perspective on Transcription Factor Binding in Arabidopsis Thaliana.” Plant Cell 26 (10): 3894–3910. https://doi.org/10.1105/tpc.114.130591.

Hinrichs, A. S., D. Karolchik, R. Baertsch, G. P. Barber, G. Bejerano, H. Clawson, M. Diekhans, et al. 2006. “The UCSC Genome Browser Database: Update 2006.” Nucleic Acids Research 34 (suppl_1): D590–98. https://doi.org/10.1093/nar/gkj144.

Huang, Ji, Juefei Zheng, Hui Yuan, and Karen McGinnis. 2018. “Distinct Tissue-Specific Transcriptional Regulation Revealed by Gene Regulatory Networks in Maize.” BMC Plant Biology 18 (1). https://doi.org/10.1186/s12870-018-1329-y.

Hufford, Matthew B., Arun S. Seetharam, Margaret R. Woodhouse, Kapeel M. Chougule, Shujun Ou, Jianing Liu, William A. Ricci, et al. 2021. “De Novo Assembly, Annotation, and Comparative Analysis of 26 Diverse Maize Genomes.” Science 373 (6555): 655–62. https://doi.org/10.1126/science.abg5289.

Huynh-Thu, Vân Anh, Alexandre Irrthum, Louis Wehenkel, and Pierre Geurts. 2010. “Inferring Regulatory Networks from Expression Data Using Tree-Based Methods.” PLoS ONE 5 (9): 1–10. https://doi.org/10.1371/journal.pone.0012776.

Javelle, Marie, Catherine Klein-Cosson, Vanessa Vernoud, Véronique Boltz, Chris Maher, Marja Timmermans, Nathalie Depège-Fargeix, and Peter M. Rogowsky. 2011. “Genome-Wide Characterization of the HD-ZIP IV Transcription Factor Family in Maize: Preferential Expression in the Epidermis1[C][W].” Plant Physiology 157 (2): 790–803. https://doi.org/10.1104/pp.111.182147.

Javelle, Marie, Vanessa Vernoud, Nathalie Depège-Fargeix, Christine Arnould, Delphine Oursel, Frédéric Domergue, Xavier Sarda, and Peter M. Rogowsky. 2010. “Overexpression of the Epidermis-Specific Homeodomain-Leucine Zipper IV Transcription Factor OUTER CELL LAYER1 in Maize Identifies Target Genes Involved in Lipid Metabolism and Cuticle Biosynthesis.” Plant Physiology 154 (1): 273–86. https://doi.org/10.1104/pp.109.150540.

Jiang, Junyao, Pin Lyu, Jinlian Li, Sunan Huang, Jiawang Tao, Seth Blackshaw, Jiang Qian, and Jie Wang. 2022. “IReNA: Integrated Regulatory Network Analysis of Single-Cell Transcriptomes and Chromatin Accessibility Profiles.” IScience 25 (11). https://doi.org/10.1016/j.isci.2022.105359.

Jiao, Yinping, Paul Peluso, Jinghua Shi, Tiffany Liang, Michelle C. Stitzer, Bo Wang, Michael S. Campbell, et al. 2017. “Improved Maize Reference Genome with Single-Molecule Technologies.” Nature 546 (7659): 524–27. https://doi.org/10.1038/nature22971.

Jin, Jinpu, Feng Tian, De-Chang Yang, Yu-Qi Meng, Lei Kong, Jingchu Luo, and Ge Gao. 2017. “PlantTFDB 4.0: Toward a Central Hub for Transcription Factors and Regulatory Interactions in Plants.” Nucleic Acids Research 45 (D1): D1040–45. https://doi.org/10.1093/nar/gkw982.

Johnson, David S., Ali Mortazavi, Richard M. Myers, and Barbara Wold. 2007. “Genome-Wide Mapping of in Vivo Protein-DNA Interactions.” Science 316 (5830): 1497–1502. https://doi.org/10.1126/science.1141319.

Jones, D. Marc, and Klaas Vandepoele. 2020. “Identification and Evolution of Gene Regulatory Networks: Insights from Comparative Studies in Plants.” Current Opinion in Plant Biology 54: 42–48. https://doi.org/10.1016/j.pbi.2019.12.008.

Kajala, Kaisa, Mona Gouran, Lidor Shaar-Moshe, G. Alex Mason, Joel Rodriguez-Medina, Dorota Kawa, Germain Pauluzzi, et al. 2021. “Innovation, Conservation, and Repurposing of Gene Function in Root Cell Type Development.” Cell 184 (12): 3333–3348.e19. https://doi.org/10.1016/j.cell.2021.04.024.

Kamimoto, Kenji, Blerta Stringa, Christy M. Hoffmann, Kunal Jindal, Lilianna Solnica-Krezel, and Samantha A. Morris. 2023. “Dissecting Cell Identity via Network Inference and in Silico Gene Perturbation.” Nature 614 (7949): 742–51. https://doi.org/10.1038/s41586-022-05688-9.

Kim, Hyo Jung, Hong Gil Nam, and Pyung Ok Lim. 2016. “Regulatory Network of NAC Transcription Factors in Leaf Senescence.” Current Opinion in Plant Biology, SI: 33: Cell signalling and gene regulation 2016, 33 (October): 48–56. https://doi.org/10.1016/j.pbi.2016.06.002.

Korhonen, Janne, Petri Martinmäki, Cinzia Pizzi, Pasi Rastas, and Esko Ukkonen. 2009. “MOODS: Fast Search for Position Weight Matrix Matches in DNA Sequences.” Bioinformatics 25 (23): 3181–82. https://doi.org/10.1093/bioinformatics/btp554.

Krouk, Gabriel, Jesse Lingeman, Amy Marshall Colon, Gloria Coruzzi, and Dennis Shasha. 2013. “Gene Regulatory Networks in Plants: Learning Causality from Time and Perturbation.” Genome Biology 14 (6): 123. https://doi.org/10.1186/gb-2013-14-6-123.

Kulkarni, Shubhada R., D. Marc Jones, and Klaas Vandepoele. 2019. “Enhanced Maps of Transcription Factor Binding Sites Improve Regulatory Networks Learned from Accessible Chromatin DatA.” Plant Physiology 181 (2): 412–25. https://doi.org/10.1104/pp.19.00605.

Kulkarni, Shubhada R., and Klaas Vandepoele. 2020. “Inference of Plant Gene Regulatory Networks Using Data-Driven Methods: A Practical Overview.” Biochimica et Biophysica Acta - Gene Regulatory Mechanisms 1863 (6): 194447. https://doi.org/10.1016/j.bbagrm.2019.194447.

Kulkarni, Shubhada R., Dries Vaneechoutte, Jan Van de Velde, and Klaas Vandepoele. 2018. “TF2Network: Predicting Transcription Factor Regulators and Gene Regulatory Networks in Arabidopsis Using Publicly Available Binding Site Information.” Nucleic Acids Research 46 (6): e31. https://doi.org/10.1093/nar/gkx1279.

Lambert, Samuel A., Ally W.H. Yang, Alexander Sasse, Gwendolyn Cowley, Mihai Albu, Mark X. Caddick, Quaid D. Morris, Matthew T. Weirauch, and Timothy R. Hughes. 2019. “Similarity Regression Predicts Evolution of Transcription Factor Sequence Specificity.” Nature Genetics 51 (6): 981–89. https://doi.org/10.1038/s41588-019-0411-1.

Langfelder, Peter, and Steve Horvath. 2008. “WGCNA: An R Package for Weighted Correlation Network Analysis.” BMC Bioinformatics 9 (1): 559. https://doi.org/10.1186/1471-2105-9-559.

Lawson, Tracy, and Jack Matthews. 2020. “Guard Cell Metabolism and Stomatal Function.” Annual Review of Plant Biology 71 (1): 273–302. https://doi.org/10.1146/annurev-arplant-050718-100251.

Layer, Ryan M., Brent S. Pedersen, Tonya Disera, Gabor T. Marth, Jason Gertz, and Aaron R. Quinlan. 2018. “GIGGLE: A Search Engine for Large-Scale Integrated Genome Analysis.” Nature Methods 15 (2): 123–26. https://doi.org/10.1038/nmeth.4556.

Li, En, Han Liu, Liangliang Huang, Xiangbo Zhang, Xiaomei Dong, Weibin Song, Haiming Zhao, and Jinsheng Lai. 2019. “Long-Range Interactions between Proximal and Distal Regulatory Regions in Maize.” Nature Communications 10 (1). https://doi.org/10.1038/s41467-019-10603-4.

Li, Guoliang, Melissa J. Fullwood, Han Xu, Fabianus Hendriyan Mulawadi, Stoyan Velkov, Vinsensius Vega, Pramila Nuwantha Ariyaratne, et al. 2010. “ChIA-PET Tool for Comprehensive Chromatin Interaction Analysis with Paired-End Tag Sequencing.” Genome Biology 11 (2): R22. https://doi.org/10.1186/gb-2010-11-2-r22.

Li, Heng, Bob Handsaker, Alec Wysoker, Tim Fennell, Jue Ruan, Nils Homer, Gabor Marth, Goncalo Abecasis, Richard Durbin, and 1000 Genome Project Data Processing Subgroup. 2009. “The Sequence Alignment/Map Format and SAMtools.” Bioinformatics 25 (16): 2078–79. https://doi.org/10.1093/bioinformatics/btp352.

Li, Yihao, Xin Zhang, Yi Zhang, and Haiyun Ren. 2022. “Controlling the Gate: The Functions of the Cytoskeleton in Stomatal Movement.” Frontiers in Plant Science 13. https://www.frontiersin.org/articles/10.3389/fpls.2022.849729.

Liang, Zhikai, Zachary A. Myers, Dominic Petrella, Julia Engelhorn, Thomas Hartwig, and Nathan M. Springer. 2022. “Mapping Responsive Genomic Elements to Heat Stress in a Maize Diversity Panel.” Genome Biology 23 (1): 234. https://doi.org/10.1186/s13059-022-02807-7.

Lu, Zefu, Brigitte T. Hofmeister, Christopher Vollmers, Rebecca M. DuBois, and Robert J. Schmitz. 2017. “Combining ATAC-Seq with Nuclei Sorting for Discovery of Cis-Regulatory Regions in Plant Genomes.” Nucleic Acids Research 45 (6): e41. https://doi.org/10.1093/nar/gkw1179.

Lu, Zefu, Alexandre P. Marand, William A. Ricci, Christina L. Ethridge, Xiaoyu Zhang, and Robert J. Schmitz. 2019. “The Prevalence, Evolution and Chromatin Signatures of Plant Regulatory Elements.” Nature Plants 5 (12): 1250–59. https://doi.org/10.1038/s41477-019-0548-z.

Majeran, Wojciech, Yang Cai, Qi Sun, and Klaas J. Van Wijk. 2005. “Functional Differentiation of Bundle Sheath and Mesophyll Maize Chloroplasts Determined by Comparative Proteomics.” The Plant Cell 17 (11): 3111–40. https://doi.org/10.1105/TPC.105.035519.

Marand, Alexandre P, Zongliang Chen, Andrea Gallavotti, and Robert J Schmitz. 2021. “A Cis-Regulatory Atlas in Maize at Single-Cell Resolution.” Cell 184 (11): 3041–3055.e21. https://doi.org/10.1016/j.cell.2021.04.014.

Marbach, Daniel, James C. Costello, Robert Küffner, Nicole M. Vega, Robert J. Prill, Diogo M. Camacho, Kyle R. Allison, Manolis Kellis, James J. Collins, and Gustavo Stolovitzky. 2012. “Wisdom of Crowds for Robust Gene Network Inference.” Nature Methods 9 (8): 796–804. https://doi.org/10.1038/nmeth.2016.

Marbach, Daniel, Sushmita Roy, Ferhat Ay, Patrick E. Meyer, Rogerio Candeias, Tamer Kahveci, Christopher A. Bristow, and Manolis Kellis. 2012. “Predictive Regulatory Models in Drosophila Melanogaster by Integrative Inference of Transcriptional Networks.” Genome Research 22 (7): 1334–49. https://doi.org/10.1101/gr.127191.111.

Mariani, Luca, Kathryn Weinand, Stephen S. Gisselbrecht, and Martha L. Bulyk. 2020. “MedeA: Analysis of Transcription Factor Binding Motifs in Accessible Chromatin.” Genome Research 30 (5): 736–48. https://doi.org/10.1101/gr.260877.120.

McCalla, Sunnie Grace, Alireza Fotuhi Siahpirani, Jiaxin Li, Saptarshi Pyne, Matthew Stone, Viswesh Periyasamy, Junha Shin, and Sushmita Roy. 2023. “Identifying Strengths and Weaknesses of Methods for Computational Network Inference from Single-Cell RNA-Seq Data.” G3 Genes|Genomes|Genetics 13 (3): jkad004. https://doi.org/10.1093/g3journal/jkad004.

Mejia-Guerra, Maria Katherine, Marcelo Pomeranz, Kengo Morohashi, and Erich Grotewold. 2012. “From Plant Gene Regulatory Grids to Network Dynamics.” Biochimica et Biophysica Acta 1819 (5): 454–65. https://doi.org/10.1016/J.BBAGRM.2012.02.016.

Meng, Xiangdong, Michael H. Brodsky, and Scot A. Wolfe. 2005. “A Bacterial One-Hybrid System for Determining the DNA-Binding Specificity of Transcription Factors.” Nature Biotechnology 23 (8): 988–94. https://doi.org/10.1038/nbt1120.

Mertz, Rachel A., and Thomas P. Brutnell. 2014. “Bundle Sheath Suberization in Grass Leaves: Multiple Barriers to Characterization.” Journal of Experimental Botany 65 (13): 3371–80. https://doi.org/10.1093/JXB/ERU108.

Misra, Biswapriya B., Biswa R. Acharya, David Granot, Sarah M. Assmann, and Sixue Chen. 2015. “The Guard Cell Metabolome: Functions in Stomatal Movement and Global Food Security.” Frontiers in Plant Science 6. https://www.frontiersin.org/articles/10.3389/fpls.2015.00334.

Negi, Juntaro, Kosuke Moriwaki, Mineko Konishi, Ryusuke Yokoyama, Toshiaki Nakano, Kensuke Kusumi, Mimi Hashimoto-Sugimoto, et al. 2013. “A Dof Transcription Factor, SCAP1, Is Essential for the Development of Functional Stomata in Arabidopsis.” Current Biology 23 (6): 479–84. https://doi.org/10.1016/j.cub.2013.02.001.

Nelissen, Hilde, Bart Rymen, Yusuke Jikumaru, Kirin Demuynck, Mieke Van Lijsebettens, Yuji Kamiya, Dirk Inzé, and Gerrit T. S. Beemster. 2012. “A Local Maximum in Gibberellin Levels Regulates Maize Leaf Growth by Spatial Control of Cell Division.” Current Biology 22 (13): 1183–87. https://doi.org/10.1016/j.cub.2012.04.065.

Noshay, Jaclyn M, Alexandre P Marand, Sarah N Anderson, Peng Zhou, Maria Katherine Mejia Guerra, Zefu Lu, Christine H O’Connor, et al. 2021. “Assessing the Regulatory Potential of Transposable Elements Using Chromatin Accessibility Profiles of Maize Transposons.” Edited by K Bomblies. Genetics 217 (1): 1–13. https://doi.org/10.1093/genetics/iyaa003.

O’Malley, Ronan C., Shao shan Carol Huang, Liang Song, Mathew G. Lewsey, Anna Bartlett, Joseph R. Nery, Mary Galli, Andrea Gallavotti, and Joseph R. Ecker. 2016. “Erratum: Cistrome and Epicistrome Features Shape the Regulatory DNA Landscape (Cell (2016) 165(5) (1280–1292)).” Cell 166 (6): 1598. https://doi.org/10.1016/j.cell.2016.08.063.

Peng, Yong, Dan Xiong, Lun Zhao, Weizhi Ouyang, Shuangqi Wang, Jun Sun, Qing Zhang, et al. 2019. “Chromatin Interaction Maps Reveal Genetic Regulation for Quantitative Traits in Maize.” Nature Communications 10 (1): 2632. https://doi.org/10.1038/s41467-019-10602-5.

Preciado, Jesus, Kevin Begcy, and Tie Liu. 2022. “The Arabidopsis HDZIP Class II Transcription Factor ABA INSENSITIVE TO GROWTH 1 Functions in Leaf Development.” Journal of Experimental Botany 73 (7): 1978–91. https://doi.org/10.1093/jxb/erab523.

Qiu, Yichun, and Claudia Köhler. 2020. “Mobility Connects: Transposable Elements Wire New Transcriptional Networks by Transferring Transcription Factor Binding Motifs.” Biochemical Society Transactions 48 (3): 1005–17. https://doi.org/10.1042/BST20190937.

Quinlan, Aaron R., and Ira M. Hall. 2010. “BEDTools: A Flexible Suite of Utilities for Comparing Genomic Features.” Bioinformatics 26 (6): 841–42. https://doi.org/10.1093/bioinformatics/btq033.

Ramachandran, Vasagi, Yuki Tobimatsu, Yamamura Masaomi, Ryosuke Sano, Toshiaki Umezawa, Taku Demura, and Misato Ohtani. 2020. “Plant-Specific Dof Transcription Factors VASCULAR-RELATED DOF1 and VASCULAR-RELATED DOF2 Regulate Vascular Cell Differentiation and Lignin Biosynthesis in Arabidopsis.” Plant Molecular Biology 104 (3): 263–81. https://doi.org/10.1007/s11103-020-01040-9.

RB, Deal, and Henikoff S. 2011. “The INTACT Method for Cell Type-Specific Gene Expression and Chromatin Profiling in Arabidopsis Thaliana.” Nature Protocols 6 (1): 56–68. https://doi.org/10.1038/NPROT.2010.175.

Reynoso, Mauricio A., Kaisa Kajala, Marko Bajic, Donnelly A. West, Germain Pauluzzi, Andrew I. Yao, Kathryn Hatch, et al. 2019. “Evolutionary Flexibility in Flooding Response Circuitry in Angiosperms.” Science 365 (6459): 1291–95. https://doi.org/10.1126/science.aax8862.

Riaño-Pachón, Diego Mauricio, Slobodan Ruzicic, Ingo Dreyer, and Bernd Mueller-Roeber. 2007. “PlnTFDB: An Integrative Plant Transcription Factor Database.” BMC Bioinformatics 8 (1): 42. https://doi.org/10.1186/1471-2105-8-42.

Roy, Sushmita, Stephen Lagree, Zhonggang Hou, James A. Thomson, Ron Stewart, and Audrey P. Gasch. 2013. “Integrated Module and Gene-Specific Regulatory Inference Implicates Upstream Signaling Networks.” PLOS Computational Biology 9 (10): e1003252. https://doi.org/10.1371/journal.pcbi.1003252.

Saelens, Wouter, Robrecht Cannoodt, and Yvan Saeys. 2018. “A Comprehensive Evaluation of Module Detection Methods for Gene Expression Data.” Nature Communications 9 (1). https://doi.org/10.1038/s41467-018-03424-4.

Savadel, Savannah D., Thomas Hartwig, Zachary M. Turpin, Daniel L. Vera, Pei-Yau Lung, Xin Sui, Max Blank, et al. 2021. “The Native Cistrome and Sequence Motif Families of the Maize Ear.” Edited by Nathan M. Springer. PLOS Genetics 17 (8): e1009689. https://doi.org/10.1371/journal.pgen.1009689.

Schmitz, Robert J., Erich Grotewold, and Maike Stam. 2022. “Cis-Regulatory Sequences in Plants: Their Importance, Discovery, and Future Challenges.” The Plant Cell 34 (2): 718–41. https://doi.org/10.1093/PLCELL/KOAB281.

Seok, Hye-Yeon, Huong Thi Tran, Sun-Young Lee, and Yong-Hwan Moon. 2022. “AtERF71/HRE2, an Arabidopsis AP2/ERF Transcription Factor Gene, Contains Both Positive and Negative Cis-Regulatory Elements in Its Promoter Region Involved in Hypoxia and Salt Stress Responses.” International Journal of Molecular Sciences 23 (10): 5310. https://doi.org/10.3390/ijms23105310.

Sijacic, Paja, Marko Bajic, Elizabeth C. McKinney, Richard B. Meagher, and Roger B. Deal. 2018. “Changes in Chromatin Accessibility between Arabidopsis Stem Cells and Mesophyll Cells Illuminate Cell Type-Specific Transcription Factor Networks.” Plant Journal 94 (2): 215–31. https://doi.org/10.1111/tpj.13882.

Simmons, Abigail R., Kelli A. Davies, Wanpeng Wang, Zhongchi Liu, and Dominique C. Bergmann. 2019. “SOL1 and SOL2 Regulate Fate Transition and Cell Divisions in the Arabidopsis Stomatal Lineage.” Development 146 (3): dev171066. https://doi.org/10.1242/dev.171066.

Song, Liang, Shao Shan Carol Huang, Aaron Wise, Rosa Castanoz, Joseph R. Nery, Huaming Chen, Marina Watanabe, Jerushah Thomas, Ziv Bar-Joseph, and Joseph R. Ecker. 2016. “A Transcription Factor Hierarchy Defines an Environmental Stress Response Network.” Science 354 (6312). https://doi.org/10.1126/SCIENCE.AAG1550.

Strable, Josh, and Hilde Nelissen. 2021. “The Dynamics of Maize Leaf Development: Patterned to Grow While Growing a Pattern.” Current Opinion in Plant Biology, Cell signaling and gene regulation, 63 (October): 102038. https://doi.org/10.1016/j.pbi.2021.102038.

Sturm, Marc, Christopher Schroeder, and Peter Bauer. 2016. “SeqPurge: Highly-Sensitive Adapter Trimming for Paired-End NGS Data.” BMC Bioinformatics 17 (1): 208. https://doi.org/10.1186/s12859-016-1069-7.

Suhita, Dontamala, Agepati S. Raghavendra, June M. Kwak, and Alain Vavasseur. 2004. “Cytoplasmic Alkalization Precedes Reactive Oxygen Species Production during Methyl Jasmonate- and Abscisic Acid-Induced Stomatal Closure.” Plant Physiology 134 (4): 1536–45. https://doi.org/10.1104/pp.103.032250.

Tello-Ruiz, Marcela Karey, Pankaj Jaiswal, and Doreen Ware. 2022. “Gramene: A Resource for Comparative Analysis of Plants Genomes and Pathways.” Methods in Molecular Biology (Clifton, N.J.) 2443: 101–31. https://doi.org/10.1007/978-1-0716-2067-0_5.

Tian, Hao, Yuru Li, Ce Wang, Xingwen Xu, Yajie Zhang, Qudsia Zeb, Johan Zicola, et al. 2021. “Photoperiod-Responsive Changes in Chromatin Accessibility in Phloem-Companion and Epidermis Cells of Arabidopsis Leaves.” The Plant Cell, January. https://doi.org/10.1093/plcell/koaa043.

Tu, Xiaoyu, María Katherine Mejía-Guerra, Jose A Valdes Franco, David Tzeng, Po Yu Chu, Wei Shen, Yingying Wei, et al. 2020. “Reconstructing the Maize Leaf Regulatory Network Using ChIP-Seq Data of 104 Transcription Factors.” Nature Communications 11 (1). https://doi.org/10.1038/s41467-020-18832-8.

Van Bel, Michiel, Tim Diels, Emmelien Vancaester, Lukasz Kreft, Alexander Botzki, Yves Van De Peer, Frederik Coppens, and Klaas Vandepoele. 2018. “PLAZA 4.0: An Integrative Resource for Functional, Evolutionary and Comparative Plant Genomics.” Nucleic Acids Research 46 (D1): D1190–96. https://doi.org/10.1093/nar/gkx1002.

Vercruysse, Jasmien, Alexandra Baekelandt, Nathalie Gonzalez, and Dirk Inzé. 2021. “Molecular Networks Regulating Cell Division during Arabidopsis Leaf Growth.” Journal of Experimental Botany 71 (8): 2365–78. https://doi.org/10.1093/JXB/ERZ522.

Wang, Xiao’e, B. M. Vindhya S. Basnayake, Huijuan Zhang, Guojun Li, Wei Li, Nasar Virk, Tesfaye Mengiste, and Fengming Song. 2009. “The Arabidopsis ATAF1, a NAC Transcription Factor, Is a Negative Regulator of Defense Responses Against Necrotrophic Fungal and Bacterial Pathogens.” Molecular Plant-Microbe Interactions® 22 (10): 1227–38. https://doi.org/10.1094/MPMI-22-10-1227.

Weirauch, Matthew T., Ally Yang, Mihai Albu, Atina G. Cote, Alejandro Montenegro-Montero, Philipp Drewe, Hamed S. Najafabadi, et al. 2014. “Determination and Inference of Eukaryotic Transcription Factor Sequence Specificity.” Cell 158 (6): 1431–43. https://doi.org/10.1016/j.cell.2014.08.009.

Wilkins, Olivia, Christoph Hafemeister, Anne Plessis, Meisha-Marika Holloway-Phillips, Gina M. Pham, Adrienne B. Nicotra, Glenn B. Gregorio, et al. 2016. “EGRINs (Environmental Gene Regulatory Influence Networks) in Rice That Function in the Response to Water Deficit, High Temperature, and Agricultural Environments.” The Plant Cell 28 (10): 2365–84. https://doi.org/10.1105/tpc.16.00158.

Wong, Min May, Govinal Badiger Bhaskara, Tuan-Nan Wen, Wen-Dar Lin, Thao Thi Nguyen, Geeng Loo Chong, and Paul E. Verslues. 2019. “Phosphoproteomics of Arabidopsis Highly ABA-Induced1 Identifies AT-Hook–Like10 Phosphorylation Required for Stress Growth Regulation.” Proceedings of the National Academy of Sciences 116 (6): 2354–63. https://doi.org/10.1073/pnas.1819971116.

Woodhouse, Margaret R., Ethalinda K. Cannon, John L. Portwood, Lisa C. Harper, Jack M. Gardiner, Mary L. Schaeffer, and Carson M. Andorf. 2021. “A Pan-Genomic Approach to Genome Databases Using Maize as a Model System.” BMC Plant Biology 21 (1): 385. https://doi.org/10.1186/s12870-021-03173-5.

Yamaguchi, Masatoshi, Misato Ohtani, Nobutaka Mitsuda, Minoru Kubo, Masaru Ohme-Takagi, Hiroo Fukuda, and Taku Demura. 2010. “VND-INTERACTING2, a NAC Domain Transcription Factor, Negatively Regulates Xylem Vessel Formation in Arabidopsis.” The Plant Cell 22 (4): 1249–63. https://doi.org/10.1105/tpc.108.064048.

Yang, So-Dam, Pil Joon Seo, Hye-Kyung Yoon, and Chung-Mo Park. 2011. “The Arabidopsis NAC Transcription Factor VNI2 Integrates Abscisic Acid Signals into Leaf Senescence via the COR/RD Genes.” The Plant Cell 23 (6): 2155–68. https://doi.org/10.1105/tpc.111.084913.

Yuan, Guo-Cheng, Yuen-Jong Liu, Michael F. Dion, Michael D. Slack, Lani F. Wu, Steven J. Altschuler, and Oliver J. Rando. 2005. “Genome-Scale Identification of Nucleosome Positions in S. Cerevisiae.” Science 309 (5734): 626–30. https://doi.org/10.1126/science.1112178.

Zhang, Yong, Tao Liu, Clifford A. Meyer, Jérôme Eeckhoute, David S. Johnson, Bradley E. Bernstein, Chad Nusbaum, et al. 2008. “Model-Based Analysis of ChIP-Seq (MACS).” Genome Biology 9 (9): R137. https://doi.org/10.1186/gb-2008-9-9-r137.

Zhou, Peng, Zhi Li, Erika Magnusson, Fabio Gomez Cano, Peter A. Crisp, Jaclyn M. Noshay, Erich Grotewold, Candice N. Hirsch, Steven P. Briggs, and Nathan M. Springer. 2020. “Meta Gene Regulatory Networks in Maize Highlight Functionally Relevant Regulatory Interactions.” Plant Cell 32 (5): 1377–96. https://doi.org/10.1105/tpc.20.00080.

