## Supplementary material for "MINI-AC: Inference of plant gene regulatory networks using bulk or single-cell accessible chromatin profiles": Figure S

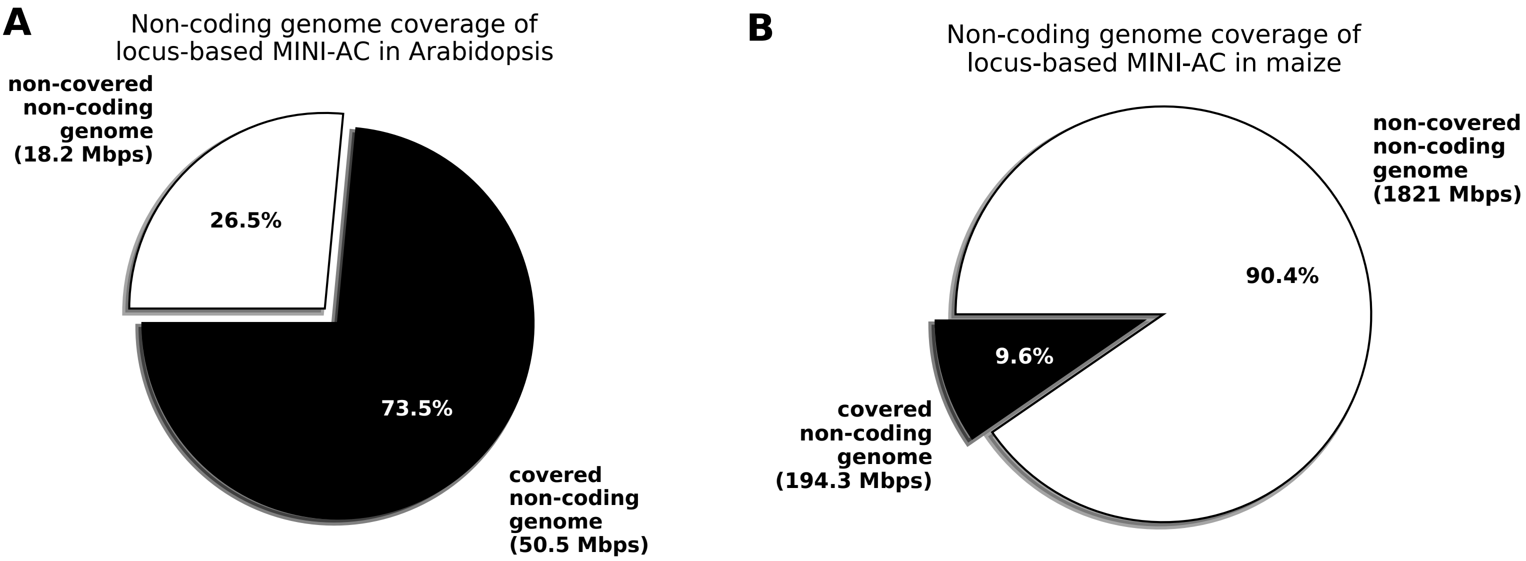


**Supplementary Figure 1. Genome coverage of Arabidopsis and maize using the non-coding genomic space extraction strategies of locus-based MINI-AC mode**. Pie charts showing the percentage of non-coding genome covered by the extraction strategy of the MINI-AC locus-based mode in Arabidopsis (A) and maize (B) using 5 kb upstream of the TrSS, introns, and 1 kb downstream of the TrES. Black represents the part of the non-coding genome that is covered, and white represents the part that is not covered. In parentheses, there is the absolute number of mega base-pairs represented by each percentage.

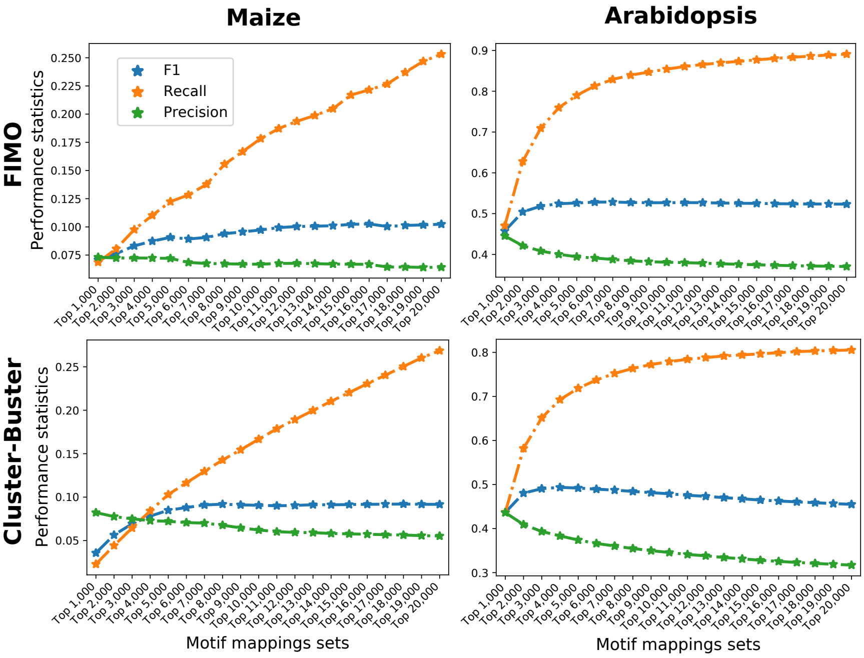


**Supplementary Figure 2. Calibration of optimal top-scoring motif matches from CB and FIMO in Arabidopsis and maize.** Performance statistics precision, recall, and F1 for different top-scoring sets of motif matches for FIMO and CB in Arabidopsis and maize by overlapping them with a ChIP-seq derived gold standard set of TFBS.

**A**

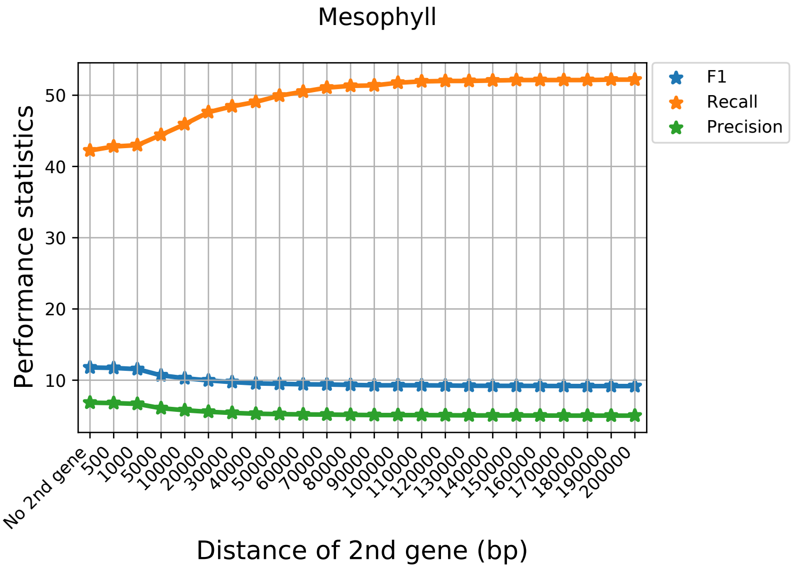


**B**

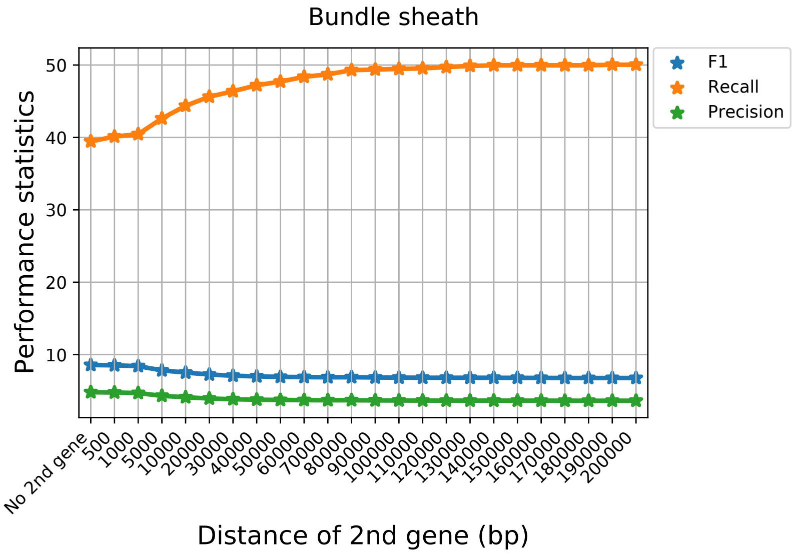


**Supplementary Figure 3. Assessment of second-closest gene annotation strategy.** Performance statistics for retrieval of cell-type specific DE genes –(A) mesophyll, (B) bundle sheath–, when annotating only the closest gene to a peak and adding progressively the second-closest genes that fall within a certain distance cutoff (x-axis).

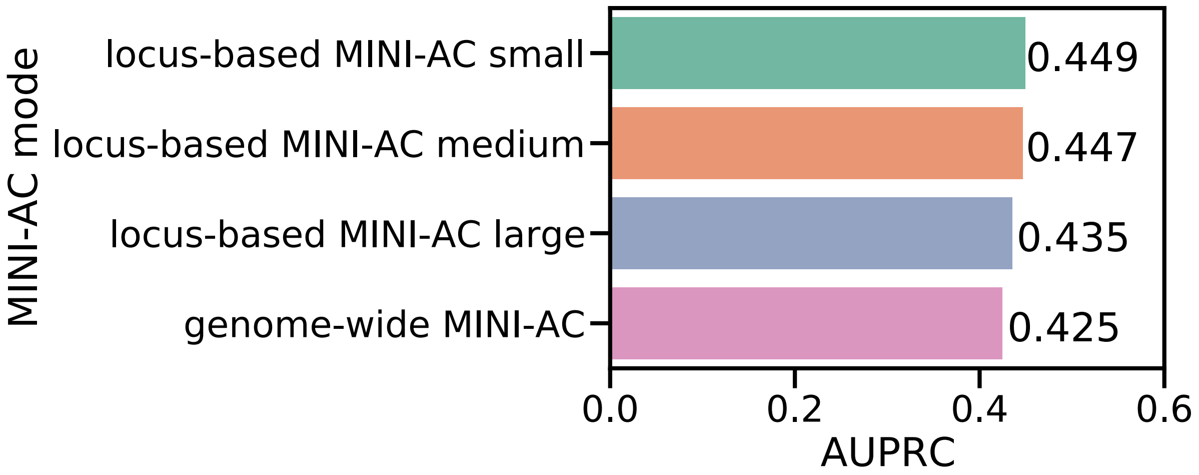


**Supplementary Figure 4. Benchmark of locus-based and genome-wide MINI-AC modes on maize.** Area under the precision-recall curve (AUPRC) for the retrieval of enriched motifs by MINI-AC that are associated with TFs of the maize leaf gold standard, for the four MINI-AC modes and non-coding genomic search spaces tested (locus-based or genome-wide).

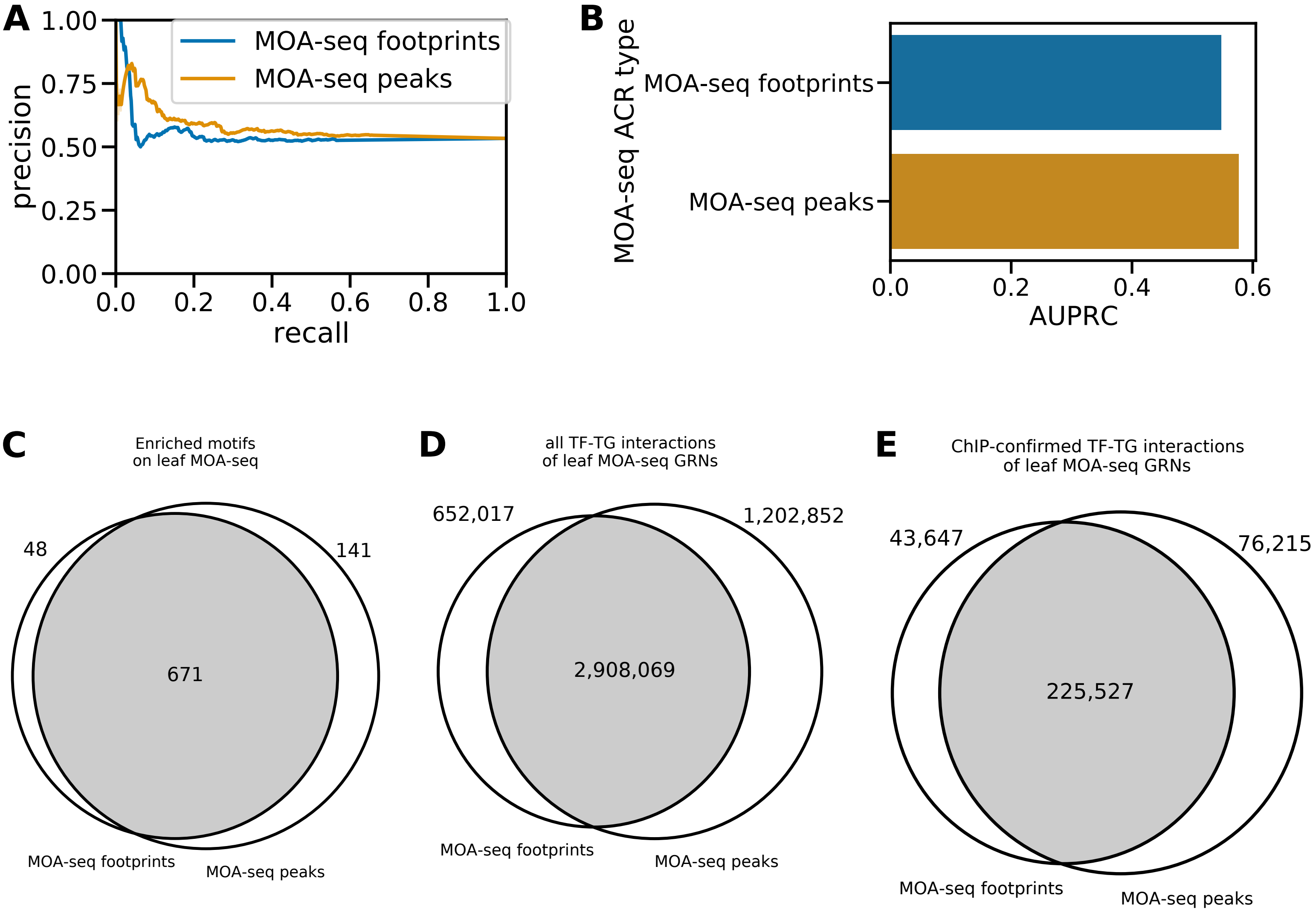


**Supplementary Figure 5. Evaluation and comparison of motif enrichment and GRNs predicted by MINI-AC using peaks and footprints derived from maize leaf MOA-seq.** (A) Precision-recall curve obtained by evaluating ranked motif enrichment results in MOA-seq peaks and footprints using a set of combined DE cell-types specific genes. (B) AUPRC relative to precision-recall curve of panel A. (C) Venn diagram showing the number of unique and shared enriched motifs predicted from MOA-seq peaks and footprints. (D) Venn diagram of the number of unique and shared TF-TG interactions of GRNs predicted using MOA-seq peaks and footprints. (E) Venn diagram of the number of unique and shared ChIP-confirmed TF-TG interactions of GRNs predicted using MOA-seq peaks and footprints.

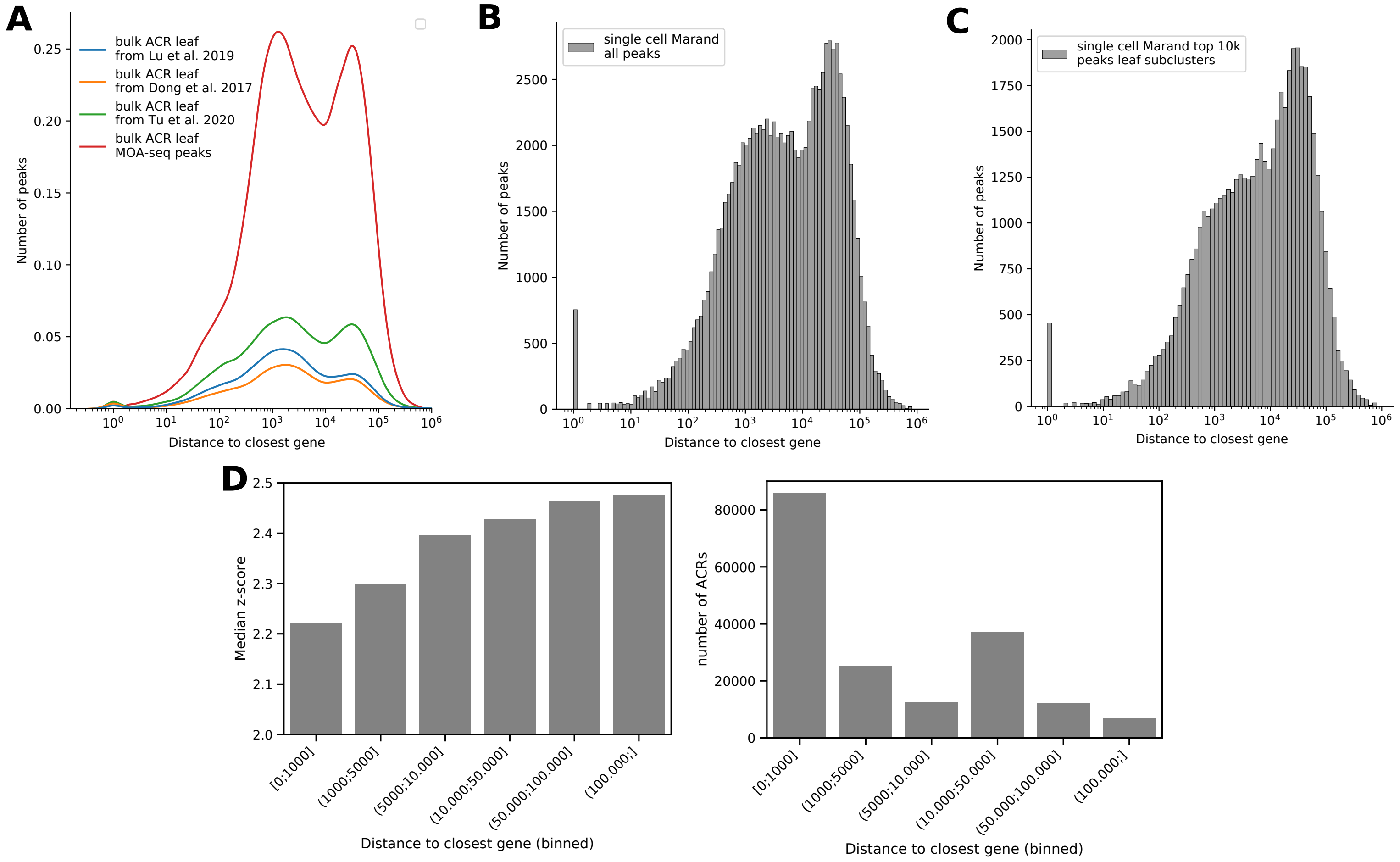


**Supplementary Figure 6. Comparison of ATAC-Seq peaks derived from bulk and single-cell technologies.** (A) Distribution plot of the distance to the closest gene for the indicated bulk ACR datasets. (B) Distribution plot of the distance to the closest gene for all the peaks of the Marand dataset. (C) Distribution plot of the distance to the closest gene for the top 10,000 z-scoring peaks of the Marand dataset. (D) Bar plot showing the median z-score (proxy for cell-type specificity) for the top 10,000 z-scoring peaks of the Marand dataset, previously binned by distance to the closest gene.

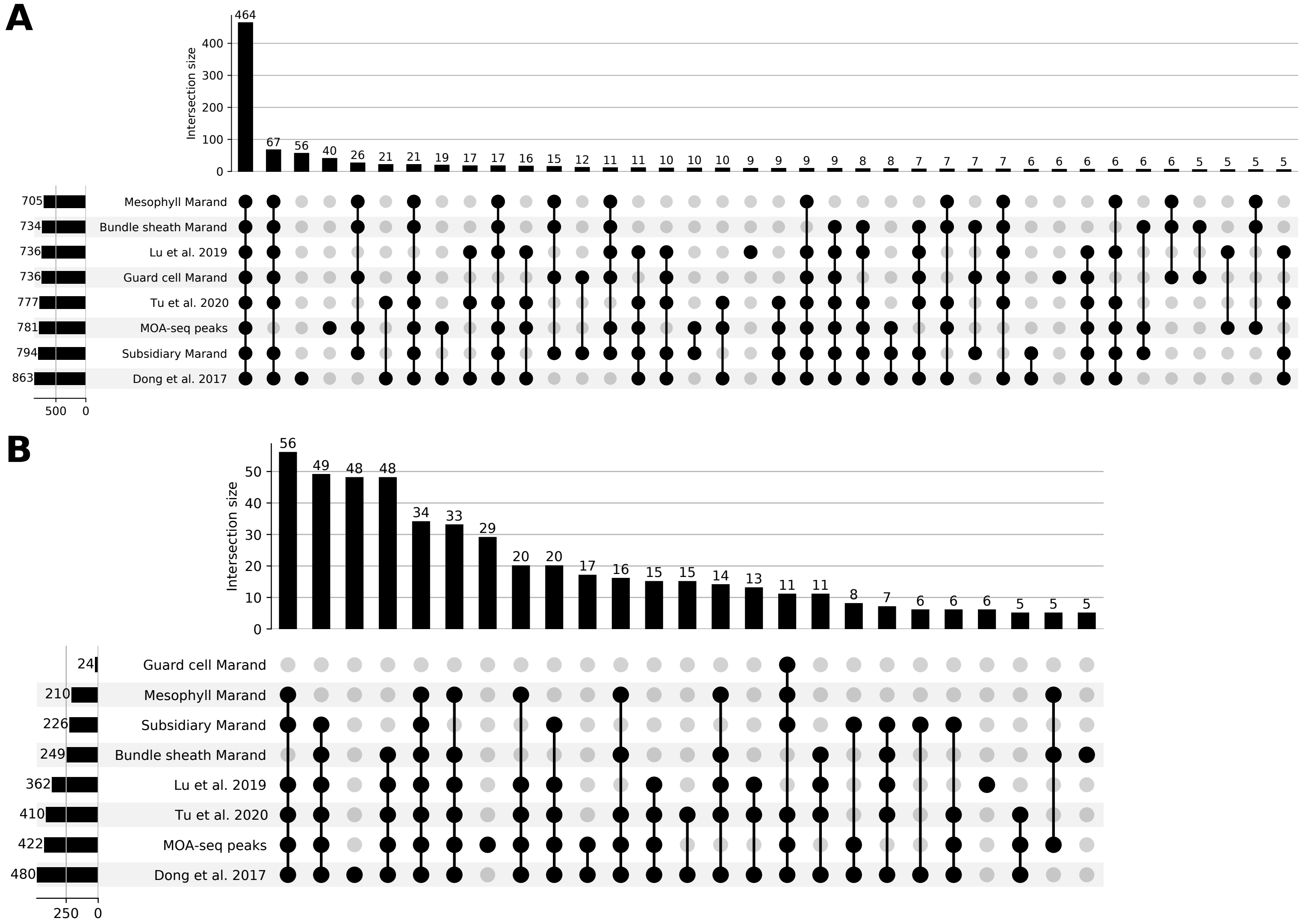


**Supplementary Figure 7. UpSet plot comparing the total number of enriched motifs and enriched motifs associated with DE TFs predicted for different bulk and single-cell leaf datasets.** (A) Comparison of the shared and unique enriched motifs for the different bulk and single-cell ACR sets. For visualization purposes, we only show overlapping sets with more than five elements. (B) Comparison of the shared and unique enriched motifs associated with DE TFs for the different bulk and single-cell ACR sets. For visualization purposes, we only show overlapping sets with more than 5 elements.

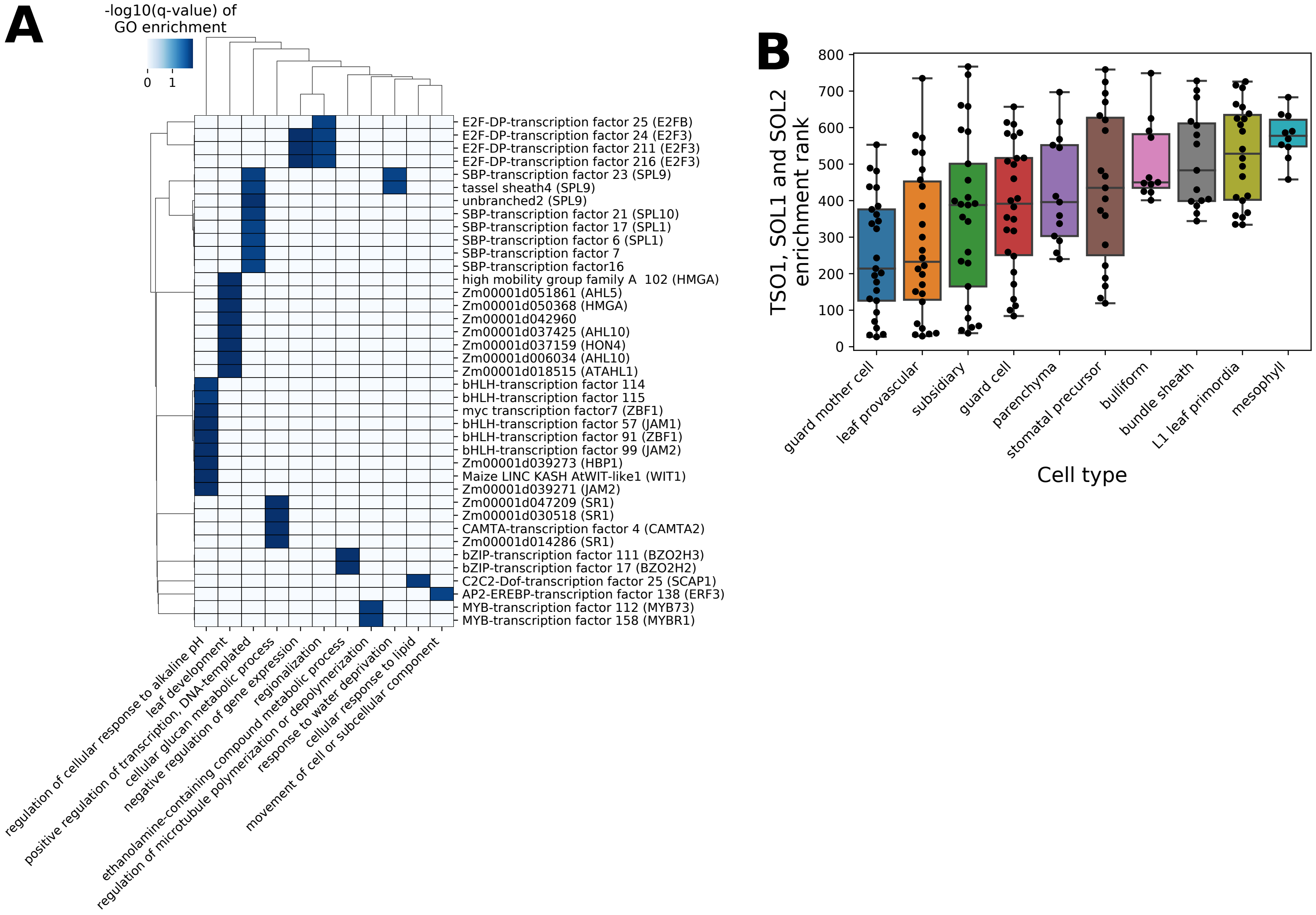


**Supplementary Figure 8. Functional network analysis of guard cells and rank distribution of TSO1 ortholog in maize.** (A) Heatmap representing the GO enrichment of the MINI-AC GRN for guard cells (all genes, DE and not DE). The GO term "cellular response to lipid" enriched for several DOF TFs, but for visualization purposes only DOF25, a maize ortholog of the Arabidopsis gene SCAP1, was kept, as SCAP1 is a known regulator of Arabidopsis guard cell function. (B) Bar plot showing the rank distribution of motifs associated with TSO1, SOL1 and SOL2 on different leaf cell types of maize.

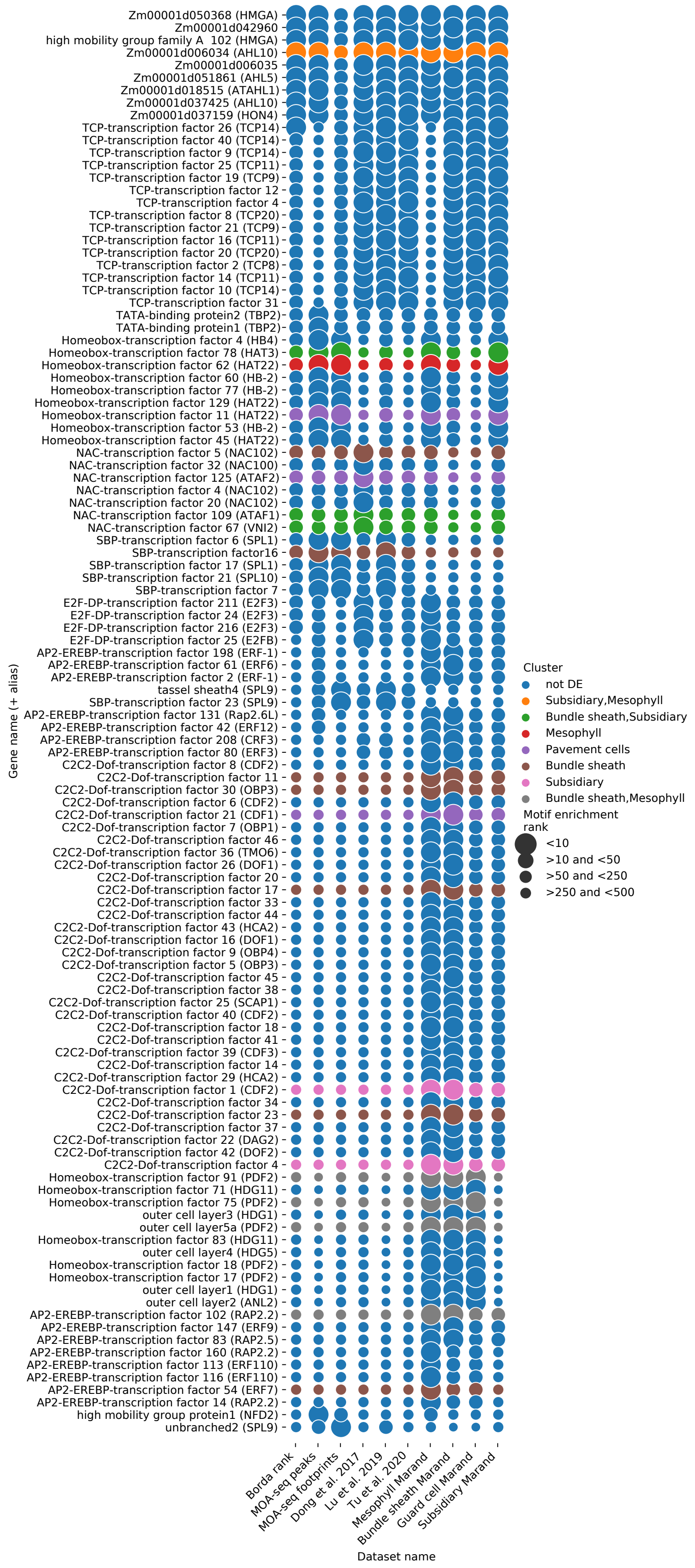


**Supplementary Figure 9. Comparison of motif enrichment ranks across different bulk and single-cell ACR sets of maize leaf.** Bubble plot showing the rank of all the TFs associated with motifs within the top 10 enrichment rank of any dataset in all the different bulk and single-cell ACR sets. The rows are colored based on the DE status of the TF on different leaf cell types, while the bubble size represents the motif enrichment rank. The lower the rank, the bigger the bubble.

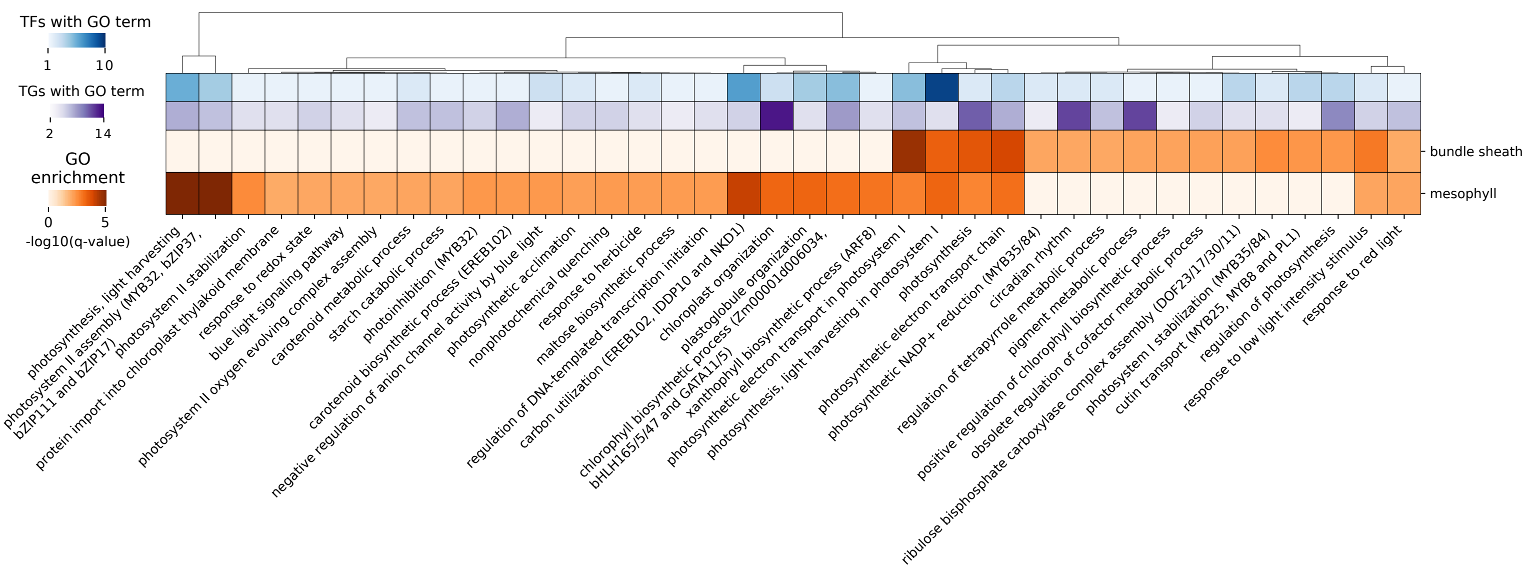


**Supplementary Figure 10. Functional analysis of maize mesophyll and bundle sheath GRNs predicted by MINI-AC.** Heatmap summarizing the GO enrichment significance (measured by -log10(q-value) in orange) of bundle sheath and mesophyll MINI-AC regulons GRNs, considering only DE TF and TGs. The row annotations are, in blue, the number of TFs with such enriched GO terms (mean if the enrichment is in the two cell types) and, in purple, the number of TGs annotated with this GO term regulated by the TF (mean if the enrichment is in the two cell types).

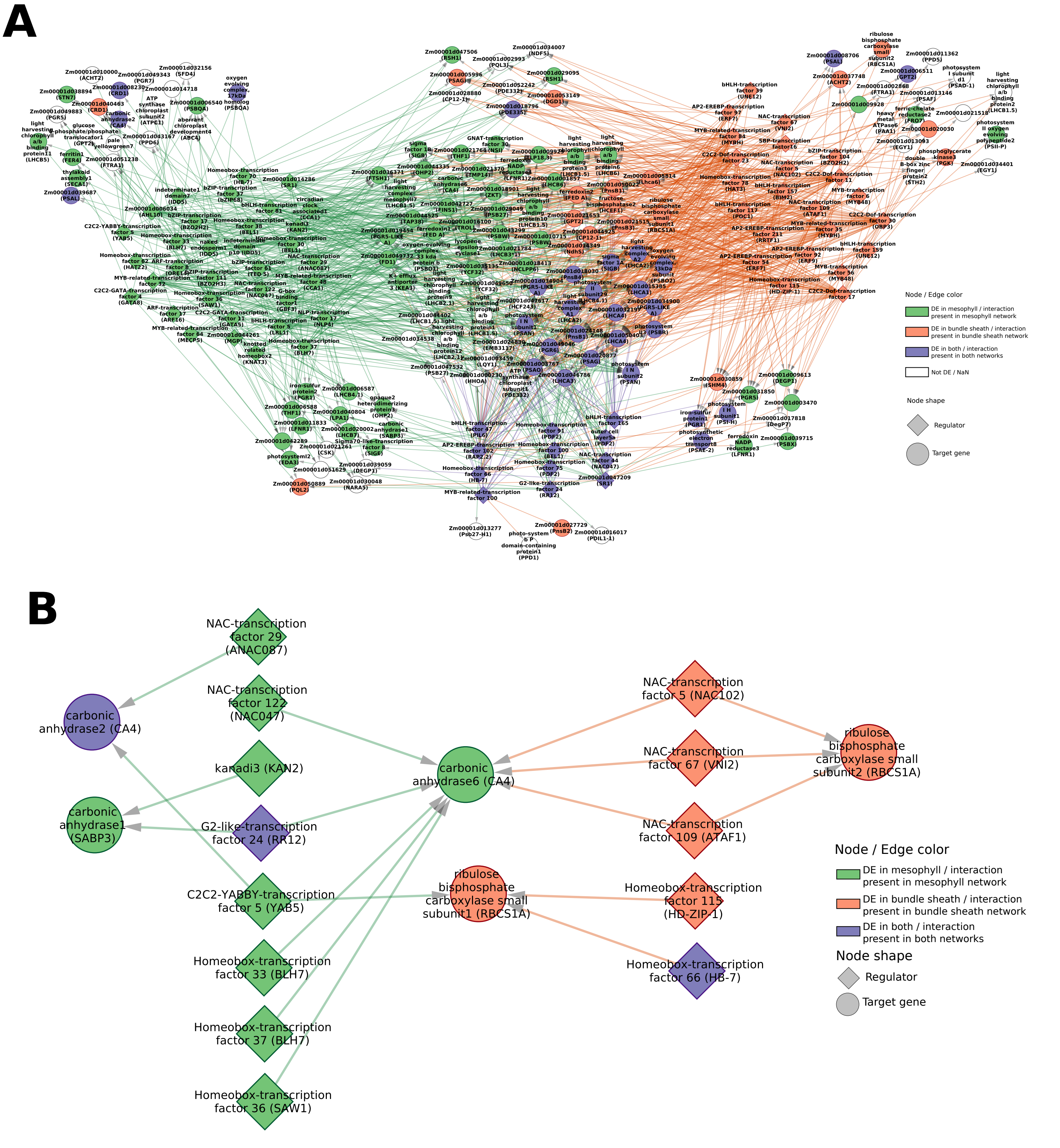


**Supplementary Figure 11. Photosynthesis GRNs of mesophyll and bundle sheath predicted by MINI-AC.** The node color indicates in which cell types the genes are DE: green for mesophyll, orange for bundle sheath, and purple for both. The edge color indicates in which GRN the interaction is present: green for mesophyll, orange for bundle sheath, and purple for both. The diamond-shaped nodes represent regulators, while the circle-shaped nodes represent target genes. (A) Mesophyll and bundle sheath GRNs predicted by MINI-AC, filtered for TFs that are DE in those specific cell types and TGs that are annotated to “photosynthesis” GO term or child terms of it. The nodes have been distributed, so the bundle sheath DE TFs are in the upper right part, the mesophyll TFs are in the upper left part, and the DE TFs in both cell types are down. The TGs controlled by the three groups of TFs are in the center. The TFs controlled uniquely by one group of TFs are in the outer part. The TGs shared between two groups of TFs are found in between the corresponding TF groups. (B) Regulators predicted by MINI-AC for carbonic anhydrase 1, 2, and 6, and RuBisCO subunits 1 and 2.
