## Supplementary material for "MINI-AC: Inference of plant gene regulatory networks using bulk or single-cell accessible chromatin profiles": Method S

### Supplementary methods 1: calibration of the optimal set of top-scoring motif mappings

The calibration of the optimal set of motif matches in the locus-based motif mappings was done individually for each motif mapping tool and each species. The motif matches were sorted by score and accumulative sets were taken from 1,000 to 20,000 in intervals of 1,000. Each set was evaluated using a gold standard of TFBS derived from ChIP-seq studies. Precision, recall, and F1 (harmonic mean of precision and recall) were computed for each set. Precision was calculated as the number of motif matches confirmed by ChIP-seq divided by the total number of matches. Recall was calculated as the number of ChIP-seq peaks for the studied TF that overlapped with motif matches, divided by the total number of ChIP-seq peaks. The set of top-scoring motif matches selected were the ones that yielded the highest F1 value for each species and motif mapping tool. For Arabidopsis, the same gold standard as in (Kulkarni, Marc Jones, and Vandepoele 2019) was used (157,363 TF binding events covering 40 TFs for which ChIP-seq data was available). For maize, the gold standard was made of ChIP-seq peaks corresponding to 62 of the 104 TFs profiled in (Tu et al. 2020) that were correctly linked to motifs present in our motif collection. The ChIP-seq peaks were downloaded from Gene Expression Omnibus (GEO); accession number GSE137972). The ChIP-seq datasets of Opaque2, RAMOSA1, Pericarp color 1, bZIP17, Knotted1 and CCA1 were added to the maize gold standard because they are associated with a motif in our collection (downloaded from GEO with accession numbers GSE63991, GSE51050, GSE38587, GSE111001, GSE39161, and GSE67655) (Bolduc et al. 2012; Morohashi et al. 2012; Eveland et al. 2014; Li et al. 2015; Ko et al. 2016; Srivastava et al. 2018). To increase the quality of the maize ChIP-seq peaks, the peak coordinates were overlapped with a merged collection of ACRs in maize coming from different studies (Oka et al. 2017; Burgess et al. 2019; Lu et al. 2019; Ricci et al. 2019) (downloaded from GEO with accession numbers GSE94291, GSE97369, GSE128434, and GSE120304). In total, there were 1,605,949 peaks belonging to 68 TFs. For each motif, motif mapping tool, and species, we determined the lowest score of calibrated top-scoring motif matches and used it as a threshold to filter the genome-wide motif mappings.

Using the same maize gold standard, the complementarity between FIMO and CB was assessed in maize. For this, motif mapping was first done using FIMO and CB on the sequences of the ChIP-seq peaks. Next, we counted the total number ChIP-seq peaks that had a FIMO or CB motif match of a PWM associated with the profiled TF, and determined the fraction of unique matches per motif mapping tool.

#### Supplementary methods 2: cell-type specific DE genes

The Arabidopsis bulk cell-type specific TFs of stem cells (59 genes) and mesophyll (158 genes) were obtained from (Kulkarni, Marc Jones, and Vandepoele 2019) (original study (Sijacic et al. 2018)). They are genes with an expression rank ratio higher than 1 (rank ratio or RR: -log2(gene expression rank in cell-type X/gene expression rank in cell-type Y)), meaning that the expression rank of these genes in one cell-type is at least two times lower than in the other cell-type. The Arabidopsis bulk cell-type specific DE genes of phloem (408 genes) and epidermis (257 genes) were obtained from the supplementary dataset 3 of (Tian et al. 2021) (induced DE genes). The single-cell derived cell-type specific DE genes for 5 maize cell types (mesophyll, bundle sheath, guard cell, subsidiary, and pavement cell) were obtained by combining data from 2 studies: one single-nuclei dataset (G. Sun et al. 2022) (supplementary dataset 6) and one single-cell dataset (Bezrutczyk et al. 2021) (supplementary dataset 1). For both datasets, a q-value of 0.05 and a log2 fold-change of 0.25 was used. The final DE genes for each cell type were: 1982 for mesophyll, 1933 for bundle sheath, 289 for guard cell, 1412 for subsidiary, and 1691 for pavement cell.

#### Supplementary methods 3: leaf-expressed genes in maize

A list of maize genes expressed in leaf was compiled by combining expression data from two different studies: one microarray experiment (Nelissen et al. 2018) and one RNA-seq experiment (X. Sun et al. 2017). For the microarray, we selected the expressed genes in the well-watered and wild-type samples. Likewise, for the RNA-seq dataset, we selected genes with CPM > 2 in the wild-type well-watered samples. To convert the AGPv3 gene IDs from these studies to AGPv4, the same gene ID mapping table as in section “Integration and curation of TF motifs” was used.

#### Literature cited

Bezrutczyk, Margaret, Nora R. Zöllner, Colin P.S. Kruse, Thomas Hartwig, Tobias Lautwein, Karl Köhrer, Wolf B. Frommer, and Ji Yun Kim. 2021. “Evidence for Phloem Loading via the Abaxial Bundle Sheath Cells in Maize Leaves.” *Plant Cell* 33 (3): 531–47. https://doi.org/10.1093/plcell/koaa055.

Bolduc, Nathalie, Alper Yilmaz, Maria Katherine Mejia-Guerra, Kengo Morohashi, Devin O’Connor, Erich Grotewold, and Sarah Hake. 2012. “Unraveling the KNOTTED1 Regulatory Network in Maize Meristems.” *Genes & Development* 26 (15): 1685–90. https://doi.org/10.1101/GAD.193433.112.

Burgess, Steven J., Ivan Reyna-Llorens, Sean R. Stevenson, Pallavi Singh, Katja Jaeger, and Julian M. Hibberd. 2019. “Genome-Wide Transcription Factor Binding in Leaves from C3 and C4 Grasses.” *The Plant Cell* 31 (10): 2297–2314. https://doi.org/10.1105/TPC.19.00078.

Eveland, Andrea L., Alexander Goldshmidt, Michael Pautler, Kengo Morohashi, Christophe Liseron-Monfils, Michael W. Lewis, Sunita Kumari, et al. 2014. “Regulatory Modules Controlling Maize Inflorescence Architecture.” *Genome Research* 24 (3): 431–43. https://doi.org/10.1101/gr.166397.113.

Ko, Dae Kwan, Dominica Rohozinski, Qingxin Song, Samuel H. Taylor, Thomas E. Juenger, Frank G. Harmon, and Z. Jeffrey Chen. 2016. “Temporal Shift of Circadian-Mediated Gene Expression and Carbon Fixation Contributes to Biomass Heterosis in Maize Hybrids.” *PLOS Genetics* 12 (7): e1006197. https://doi.org/10.1371/journal.pgen.1006197.

Kulkarni, Shubhada R., D. Marc Jones, and Klaas Vandepoele. 2019. “Enhanced Maps of Transcription Factor Binding Sites Improve Regulatory Networks Learned from Accessible Chromatin DatA.” *Plant Physiology* 181 (2): 412–25. https://doi.org/10.1104/pp.19.00605.

Li, Chaobin, Zhenyi Qiao, Weiwei Qi, Qian Wang, Yue Yuan, Xi Yang, Yuanping Tang, et al. 2015. “Genome-Wide Characterization of Cis-Acting DNA Targets Reveals the Transcriptional Regulatory Framework of Opaque2 in Maize.” *The Plant Cell* 27 (3): 532–45. https://doi.org/10.1105/tpc.114.134858.

Lu, Zefu, Alexandre P. Marand, William A. Ricci, Christina L. Ethridge, Xiaoyu Zhang, and Robert J. Schmitz. 2019. “The Prevalence, Evolution and Chromatin Signatures of Plant Regulatory Elements.” *Nature Plants* 5 (12): 1250–59. https://doi.org/10.1038/s41477-019-0548-z.

Morohashi, Kengo, María Isabel Casas, Maria Lorena Falcone Ferreyra, María Katherine Mejía-Guerra, Lucille Pourcel, Alper Yilmaz, Antje Feller, et al. 2012. “A Genome-Wide Regulatory Framework Identifies Maize Pericarp Color1 Controlled Genes.” *The Plant Cell* 24 (7): 2745–64. https://doi.org/10.1105/tpc.112.098004.

Nelissen, Hilde, Xiao-Huan Sun, Bart Rymen, Yusuke Jikumaru, Mikko Kojima, Yumiko Takebayashi, Rafael Abbeloos, et al. 2018. “The Reduction in Maize Leaf Growth under Mild Drought Affects the Transition between Cell Division and Cell Expansion and Cannot Be Restored by Elevated Gibberellic Acid Levels.” *Plant Biotechnology Journal* 16 (2): 615–27. https://doi.org/10.1111/pbi.12801.

Oka, Rurika, Johan Zicola, Blaise Weber, Sarah N. Anderson, Charlie Hodgman, Jonathan I. Gent, Jan Jaap Wesselink, et al. 2017. “Genome-Wide Mapping of Transcriptional Enhancer Candidates Using DNA and Chromatin Features in Maize.” *Genome Biology* 18 (1): 1–24. https://doi.org/10.1186/s13059-017-1273-4.

Ricci, William A., Zefu Lu, Lexiang Ji, Alexandre P. Marand, Christina L. Ethridge, Nathalie G. Murphy, Jaclyn M. Noshay, et al. 2019. “Widespread Long-Range Cis-Regulatory Elements in the Maize Genome.” *Nature Plants* 5 (12): 1237–49. https://doi.org/10.1038/s41477-019-0547-0.

Sijacic, Paja, Marko Bajic, Elizabeth C. McKinney, Richard B. Meagher, and Roger B. Deal. 2018. “Changes in Chromatin Accessibility between Arabidopsis Stem Cells and Mesophyll Cells Illuminate Cell Type-Specific Transcription Factor Networks.” *Plant Journal* 94 (2): 215–31. https://doi.org/10.1111/tpj.13882.

Srivastava, Renu, Zhaoxia Li, Giulia Russo, Jie Tang, Ran Bi, Usha Muppirala, Sivanandan Chudalayandi, et al. 2018. “Response to Persistent ER Stress in Plants: A Multiphasic Process That Transitions Cells from Prosurvival Activities to Cell Death.” *The Plant Cell* 30 (6): 1220–42. https://doi.org/10.1105/tpc.18.00153.

Sun, Guiling, Mingzhang Xia, Jieping Li, Wen Ma, Qingzeng Li, Jinjin Xie, Shenglong Bai, et al. 2022. “The Maize Single-Nucleus Transcriptome Comprehensively Describes Signaling Networks Governing Movement and Development of Grass Stomata.” *The Plant Cell* 34 (5): 1890–1911. https://doi.org/10.1093/plcell/koac047.

Sun, Xiaohuan, James Cahill, Tom Van Hautegem, Kim Feys, Clinton Whipple, Ondrej Novák, Sofie Delbare, et al. 2017. “Altered Expression of Maize PLASTOCHRON1 Enhances Biomass and Seed Yield by Extending Cell Division Duration.” *Nature Communications* 8 (1): 1–11. https://doi.org/10.1038/ncomms14752.

Tian, Hao, Yuru Li, Ce Wang, Xingwen Xu, Yajie Zhang, Qudsia Zeb, Johan Zicola, et al.

2021. “Photoperiod-Responsive Changes in Chromatin Accessibility in Phloem-Companion and Epidermis Cells of Arabidopsis Leaves.” *The Plant Cell*, January. https://doi.org/10.1093/plcell/koaa043.

Tu, Xiaoyu, María Katherine Mejía-Guerra, Jose A Valdes Franco, David Tzeng, Po Yu Chu, Wei Shen, Yingying Wei, et al. 2020. “Reconstructing the Maize Leaf Regulatory Network Using ChIP-Seq Data of 104 Transcription Factors.” *Nature Communications* 11 (1). https://doi.org/10.1038/s41467-020-18832-8.
